# Hexokinase 2 localizes to the nucleus in response to glucose limitation but does not regulate gene expression

**DOI:** 10.1101/2022.11.30.518474

**Authors:** Mitchell A. Lesko, Dakshayini G. Chandrashekarappa, Eric M. Jordahl, Katherine G. Oppenheimer, Ray Wesley Bowman, Chaowei Shang, Jacob Durrant, Martin C. Schmidt, Allyson F. O’Donnell

## Abstract

Glucose is the preferred carbon source for most eukaryotes, and the first step in its metabolism is phosphorylation to glucose-6-phosphate. This reaction is catalyzed by a family of enzymes called either hexokinases or glucokinases depending on their substrate specificity. The yeast *Saccharomyces cerevisiae* encodes three such enzymes, Hxk1, Hxk2 and Glk1. In yeast and mammals, some isoforms of this enzyme are found in the nucleus, suggesting a possible moonlighting function beyond glucose phosphorylation. In contrast to mammalian hexokinases, the yeast Hxk2 enzyme has been proposed to shuttle into the nucleus in glucose replete conditions where it reportedly moonlights as part of a glucose-repressive transcriptional complex. To achieve this role in glucose repression, Hxk2 reportedly binds the Mig1 transcriptional repressor, is dephosphorylated at serine 15 in its N-terminus, and requires an N-terminal nuclear localization sequence (NLS).

In this study, we use high-resolution, quantitative, fluorescent microscopy of live cells to determine the conditions, residues, and regulatory proteins required for Hxk2 nuclear localization. In direct contradiction to previous yeast studies, our quantitative imaging demonstrates that Hxk2 is largely excluded from the nucleus under glucose replete conditions but is retained in the nucleus under glucose limiting conditions. Our data show that the Hxk2 N-terminus does not contain an NLS but instead comprises sequences necessary for nuclear exclusion and multimerization regulation. Amino acid substitutions of the phosphorylated residue, serine 15, disrupt Hxk2 dimerization but have no effect on its glucose-regulated nuclear localization. Substitution of alanine at the nearby residue, lysine 13, affects both dimerization and maintenance of nuclear exclusion under glucose replete conditions. Modeling and simulation provide insight into the molecular mechanisms of this regulation. In marked contrast to earlier studies, we find that the transcriptional repressor Mig1 and the protein kinase Snf1 have little effect on Hxk2 localization. Instead, the protein kinase Tda1 is a key regulator of Hxk2 localization.

Finally, RNAseq analyses of the yeast transcriptome further dispel the idea that Hxk2 moonlights as a transcriptional repressor, demonstrating that Hxk2 has a negligible role in transcriptional regulation in both glucose replete and limiting conditions. Taken together, our studies provide a paradigm shift for the conditions, residues, and regulators controlling Hxk2 dimerization and nuclear localization. Based on our data, the nuclear translocation of Hxk2 in yeast occurs in glucose starvation conditions, a finding that aligns well with the nuclear regulation of mammalian orthologs of this enzyme. Our findings lay the foundation for future studies of Hxk2 nuclear activity.

## Introduction

The functional complexity of many proteomes is extended by “moonlighting” proteins, a term used to describe proteins with “other jobs” in the cell [1]. Enzymes with roles in signal transduction that are independent of their catalytic activities are good examples. For instance, cytochrome c is a highly conserved protein best known as a component of the mitochondrial electron transport chain. However, when cytochrome c is released from the mitochondria, it becomes an important messenger of apoptotic signaling [2]. Glycolytic enzymes in many species have moonlighting functions distinct from their catalytic potential [3]. In this study, we examine the yeast hexokinase 2 (Hxk2), which is the enzyme that catalyzes the first step of glycolysis. The ability of Hxk2 to translocate to the nucleus suggests that Hxk2 may carry out a nuclear function distinct from its catalytic role in the cytosolic glycolysis pathway.

The budding yeast *Saccharomyces cerevisiae* express three enzymes capable of phosphorylating glucose. Any one of these is sufficient to support growth on glucose as a carbon source [4]. Two of these enzymes, Hxk1 and Hxk2, are hexokinases with broad substrate specificity that includes both glucose and fructose [5]. The third enzyme, Glk1, is a glucokinase so named for its high specificity for glucose as a substrate. The Hxk1 and Hxk2 proteins are closely related paralogs with 77% identity and 89% similarity in their amino acid sequences. Glk1 is less closely related to the hexokinases (37% identity and 53% similarity) but has a paralog named Emi2 whose function is uncertain. Recombinant Emi2 protein contains detectable glucose phosphorylating activity [6] but the presence of the *EMI2* gene is not sufficient to confer growth on glucose in triple hexokinase delete cells (*hxk1Δ hxk2Δ glk1Δ*) suggesting that Emi2 does not function as a hexokinase *in vivo* [4].

This study focuses on Hxk2, a protein proposed to have a moonlighting function in the nucleus regulating gene expression [7]. The current model for Hxk2, proposed by the Moreno lab, states that Hxk2 translocates to the nucleus in glucose rich conditions [8] in a manner dependent upon (1) binding to Mig1, a transcriptional repressor [9], (2) a nuclear localization signal (NLS) in the N-terminus of Hxk2 between lysine 6 and lysine 13 [10], and (3) dephosphorylation of Hxk2 at serine 15 [11]. Once in the nucleus, Hxk2 is proposed to be one subunit of a transcriptional repressor complex that includes the DNA binding proteins Mig1 and Mig2; the Tup1 repressor; Med8, a subunit of the Mediator complex; Reg1, a regulatory subunit of the PP1 phosphatase; and the nuclear isoform of yeast AMP-activated protein kinase (AMPK) composed of the Snf1, Snf4 and Gal83 proteins. In this model, the large complex plays a critical role in glucose repression of gene expression [7].

While the studies that support this model are cited collectively >1000 times [12–17], accumulating evidence calls it into question, and there has been no independent corroboration for any aspect of the model. For instance, large scale protein-protein association studies using TAP-purification and mass spectrometry [18,19] or global two-hybrid screens [20,21] have failed to detect Hxk2 association with any of the components of the proposed nuclear complex. Targeted studies from other labs have not provided secondary confirmation of this Hxk2-containing complex or the glucose-stimulated nuclear translocation of Hxk2. Comprehensive ChIP-Seq data fail to identify this complex at the *SUC2* promoter [22], a gene regulated by glucose-repression whose transcription is reportedly controlled by Hxk2 [8,23,24]. Furthermore, some aspects of the Hxk2 model have been openly questioned in published commentaries [25]. Finally, the idea that Hxk2 is excluded from the nucleus in glucose-starvation conditions and found in the nucleus in glucose-replete conditions in yeast is counter to many of the models for hexokinase and glucokinase regulation in other organisms [26–29]. Mammalian hexokinase isoforms II and III and glucokinase (alternatively referred to as hexokinase IV) can each be nuclear localized [26,27,30]. Most data suggest that these enzymes are nuclear localized in response to glucose limitation or stress conditions, which is the opposite of what has been reported for yeast Hxk2 [26,27,30].

Unfortunately, entwined with the model of Hxk2 nuclear-cytosolic shuttling presented above is the idea that Hxk2’s oligomeric state regulates nuclear partitioning. Rigorous biochemical analyses demonstrate that in glucose replete conditions, a balance of monomeric and dimeric Hxk2 exists but Hxk2 shifts to predominantly monomeric when glucose is restricted [31–34]. A key regulator of Hxk2 dimerization is serine 15, which is phosphorylated in glucose-starvation conditions to disrupt the dimer [31–34]. The Tda1 kinase is required for serine 15 phosphorylation, while Snf1 plays only a minor role [35]. Unlike the clear link between serine 15 phosphorylation and Hxk2 monomer-dimer balance, the proposition that serine 15 regulates Hxk2 nuclear translocation is poorly supported[25].

In this study, we use high resolution, quantitative fluorescent imaging, co-immunoprecipitation, biochemical and genetic methods to study the nuclear localization, dimerization, and nuclear function of Hxk2. Our data directly contradict essentially all aspects of the current Hxk2 nuclear localization model. Most importantly, we demonstrate that Hxk2 is excluded from the nucleus in glucose replete conditions, the very time when it is proposed to operate in a nuclear complex important for glucose repression. We find that yeast Hxk2 does enter the nucleus, but only in glucose starvation conditions, which is more in line with the starvation and stress conditions that induce a similar nuclear translocation in mammalian hexokinase isoforms [26,27]. Importantly, our imaging studies differ substantially from earlier work, which examined fluorescently tagged Hxk2 localization using yeast cells that were likely experiencing glucose starvation due to a pre-incubation in medium containing glycerol and lacking glucose [9–11,36]. In contrast to these earlier studies, we maintain the indicated glucose supply for all live-cell imaging, ensuring a more accurate representation of what happens in cells under these conditions.

Using this system, we further define *cis* and *trans* regulatory elements needed to control the starvation-induced nuclear accumulation of Hxk2. We identify Hxk2 lysine 13 as required for the glucose-regulated nuclear exclusion of Hxk2 and for dimerization. In contrast to earlier studies, mutation of serine 15 did not dramatically alter the glucose-regulated nuclear accumulation of Hxk2. However, in keeping with Kriegel lab studies, serine 15 mutants of Hxk2 altered the monomer-dimer balance in cells [31–34]. Thus, Hxk2 glucose-regulated nuclear translocation and dimerization can be uncoupled. The Tda1 kinase, and not Snf1 or Mig1, was needed for Hxk2 nuclear translocation in response to glucose starvation.

Finally, our RNAseq analyses showed that the expression of the most highly glucose-repressed genes was not affected by the deletion of the *HXK2* gene. Taken together, our data strongly refute the idea that Hxk2 moonlights as a transcriptional repressor important for glucose repression. The function of nuclear Hxk2 in glucose-starved cells remains to be defined and future work will tackle this exciting aspect of Hxk2 biology, building off the new paradigm for Hxk2 nuclear translocation established herein.

## Results

### Hxk2 nuclear shuttling

We examined the impact of glucose abundance on the nuclear propensity of the three hexokinases in *S. cerevisiae*: Hxk1, Hxk2, and Glk1. Earlier studies suggested that Hxk2 from *S. cerevisiae* and *C. albicans* is nuclear excluded in glucose starvation and partially nuclear localized in abundant glucose [11,37]. However, in these earlier Hxk2 studies, cells were incubated in glucose-free medium before imaging (glycerol-containing medium or PBS in the *S. cerevisiae* or *C. albicans* experiments, respectively), perhaps to allow for more robust nuclear staining with DAPI [11,37]. We generated functional GFP-tagged, plasmid-borne versions of these hexokinases and expressed them in cells lacking their respective endogenous genes. To examine the localization of these hexokinase-GFP fusions in glucose-replete conditions (2% glucose, “high” glucose) and after acute glucose starvation (0.05% glucose, “low” glucose), we used live-cell confocal microscopy and an mScarlet-tagged nuclear marker to quantitatively assess nuclear co-localization. In marked contrast to earlier qualitative Hxk2 localization studies [9–11,36], Hxk2 was largely cytosolic in glucose-grown cells, and a portion of Hxk2 became nuclear localized upon glucose starvation (Fig 1A).

**Figure 1.**
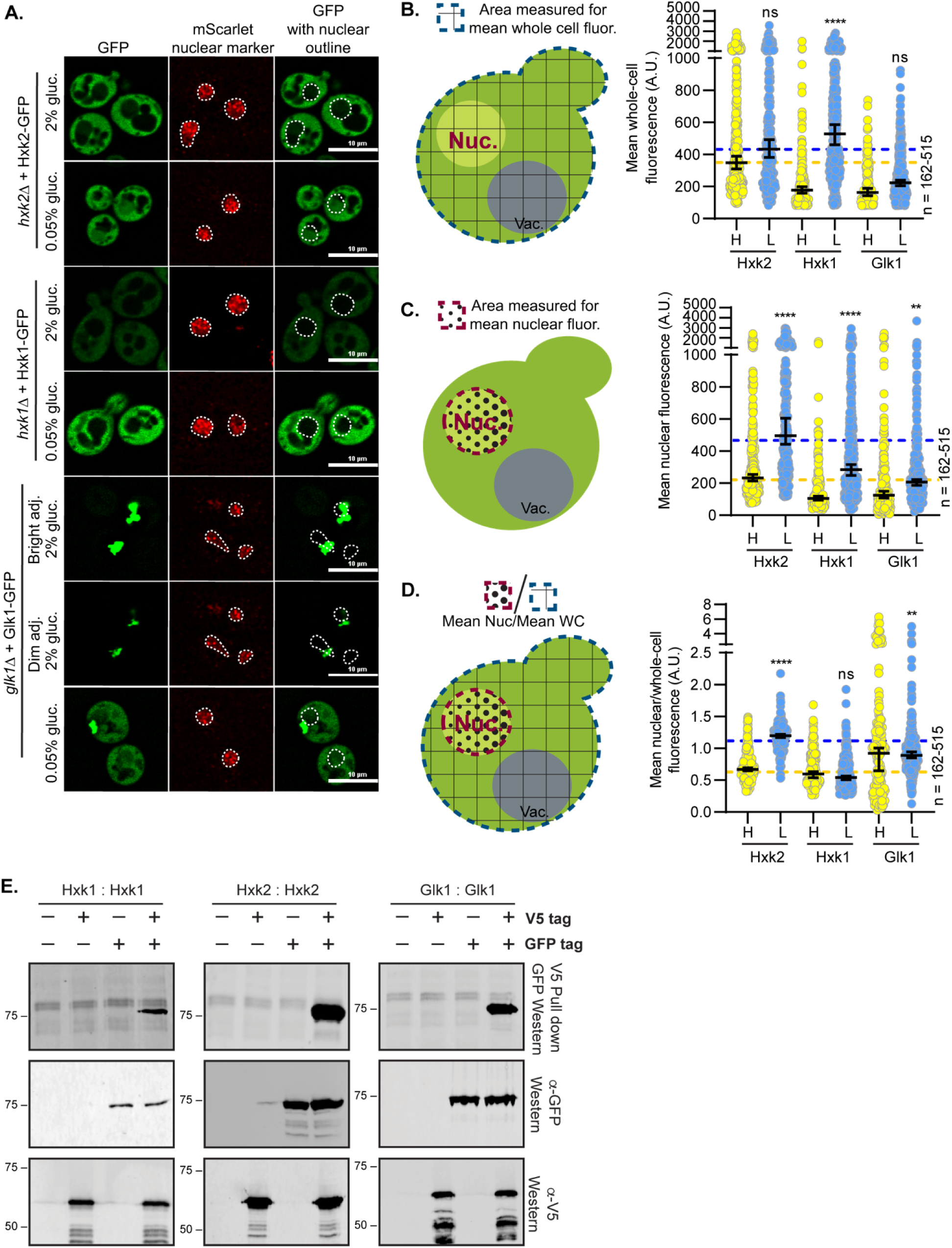
Hexokinases alter localization in response to glucose starvation and can form multimers. (A) Confocal microscopy of GFP-tagged yeast hexokinases expressed from CEN plasmids under the control of their endogenous promoters. Each hexokinase is expressed in cells where the endogenous gene has been deleted. Co-localization with the nucleus is determined based on overlap with the Tpa1-mScarlet nuclear marker, and a dashed white line indicates the nucleus. (B-D) Automated quantification (using Nikon.*ai* and GA3 analyses) of the images shown in panel A where (B) shows the mean whole-cell fluorescence intensities, (C) shows the mean nuclear fluorescence, (D) shows the mean nuclear fluorescence over the whole-cell fluorescence for each hexokinase as XY-scatter plots. The regions measured for each of these analyses are shown to the left of each graph with (B and D) the area for whole-cell measurement shown as a blue-dashed line filled with grid lines and (C-D) the area for the nuclear measurement shown as a red-dashed line filled with black dots. Horizontal bars show the median, and error bars indicate the 95% confidence interval. Dashed yellow and blue lines represent the median value for Hxk2-GFP cells in high and low glucose, respectively. Kruskal–Wallis statistical analysis with Dunn’s post hoc test was performed to compare the (B) mean whole-cell fluorescence, (C) mean nuclear fluorescence, or (D) mean nuclear/whole-cell fluorescence ratio between high and low glucose medium conditions for each hexokinase. (E) To assess multimerization, we prepared extracts from yeast cells expressing the indicated hexokinase proteins either untagged (-) or tagged (+) with V5 or GFP. Protein inputs were monitored by immunoblotting (bottom two panels). Association of tagged proteins was assessed by co-immunoprecipitation using anti-V5 beads followed by immunoblotting with anti-GFP (top panel).

To strengthen these observations, we quantified for each hexokinase: (1) the mean whole cell fluorescence, (2) mean nuclear fluorescence as a readout for nuclear abundance, and (3) the mean nuclear to mean whole-cell fluorescence ratio to assess the relative contribution of the nuclear signal to that of total cellular fluorescence (Fig 1B-D). This last measure is essential because it allows us to account for any changes in gene expression or protein abundance/stability that may accompany nutrient changes, as is evident for Hxk1 and Glk1 (Fig 1A-D). In addition, higher whole-cell fluorescence can lead to increased fluorescence in the nuclear compartment simply because the background fluorescence increases. This makes the ratio of the nuclear to whole cell fluorescence the most useful measure in assessing changes in nuclear distribution. Manual and automated image analyses were performed in four biological replicate experiments. Data from manual or automated quantification were highly consistent, validating the use of the automated quantification pipeline (Fig 1C-D and S1A-B Fig). In response to glucose starvation, the mean nuclear fluorescence of Hxk2-GFP increased 3-fold, and the ratio of nuclear-to-whole-cell Hxk2-GFP fluorescence more than doubled, demonstrating a clear shift in Hxk2 localization to the nucleus upon glucose restriction. We observed the same glucose-starvation-induced increase in nuclear fluorescence when quantitatively assessing a chromosomally integrated mNeonGreen-tagged Hxk2 using both manual and automated approaches (S1C-E Fig).

These data run counter to earlier findings that suggested Hxk2-GFP was retained in the nucleus in the presence of glucose and became nuclear excluded upon glucose starvation [9–11,36]. Three factors likely account for this discrepancy. First, as mentioned above, earlier studies incubated cells in glycerol or PBS (*i.e*., lacking glucose) before imaging [9–11,36,37]. This pre-incubation could be sufficient to provoke the glucose starvation-induced nuclear translocation we observed. Indeed, when we imaged *S. cerevisiae* after incubation in 80% glycerol, we found nuclear accumulations of Hxk2-GFP regardless of the medium in which cells were pre-grown (*i.e.,* either 2% glucose or shifted to 0.05% glucose for 2 h; S1F Fig). Second, high copy plasmids were used to express Hxk2-GFP in the earlier studies and so this overexpression might have made localization changes more difficult to interpret. Third, our studies may also differ from earlier studies due to the improvements in microscopy since the *S. cerevisiae* Hxk2-GFP images in previous studies were acquired and the lack of quantification of earlier datasets [9–11,36].

### Hxk1 and Glk1 nuclear shuttling

In contrast to Hxk2, we saw no qualitative nuclear accumulation of Hxk1 in glucose-grown or -starved cells (Fig 1A). Hxk1 transcription is upregulated upon glucose starvation, resulting in an ∼2.5-fold change in Hxk1 whole-cell fluorescence intensity (Fig 1B). If we consider only mean nuclear fluorescence, Hxk1 appears to undergo modest nuclear accumulation in glucose starvation that likely results from increased total Hxk1-GFP expression (Fig 1C). When data were normalized to account for protein abundance changes using the ratio of mean nuclear to mean whole-cell fluorescence intensity, we saw no change in the relative distribution of Hxk1 upon glucose restriction (Fig 1C-D). As expected, the abundance of Hxk1 was lower than that of Hxk2 in glucose replete conditions, but in glucose starvation Hxk1 protein levels rose to be higher than those of Hxk2 (Fig 1B). Therefore, if there was a significant accumulation of Hxk1 in the nucleus we would have certainly been able to detect it in the glucose starvation conditions, and we did not (Fig 1A-C).

Consistent with recent reports [38], Glk1 formed cytosolic inclusions in high glucose (Fig 1A). These inclusions were diminished in low-glucose conditions, where Glk1 may be mobilized to help phosphorylate glucose [38]. It is important to note that our Glk1 quantitative analyses were somewhat confounded by the large cytosolic inclusions that were often present near the nuclear marker; some cytosolic fluorescent signal was occasionally captured in the nucleus due to this overlap. Glk1 is transcriptionally upregulated in response to glucose starvation, causing a modest increase in its whole-cell fluorescence intensity (Fig1B) [39]. As with Hxk1, changes in mean nuclear fluorescence suggested Glk1 may have undergone a modest nuclear accumulation in glucose starvation, but when the data were normalized to whole-cell-fluorescence intensity, we observed no change in the relative distribution of Glk1 upon glucose restriction (Fig 1C-D). These findings suggest that, while Hxk2 undergoes a glucose starvation-induced nuclear translocation, Hxk1 and Glk1 do not.

### Hexokinase multimerization

Hxk2 nuclear localization could be linked to its transition from a dimer to a monomer. In glucose rich medium, Hxk2 exists in a balance between dimeric and monomeric species [34,35]. Upon glucose starvation, this balance shifts toward the monomeric state [34,35]. Hxk2 phosphorylation at S15 is vital for transition to the monomer. The Moreno lab suggested that in rich glucose conditions, Hxk2 is both nuclear and cytosolic but in response to glucose starvation, Hxk2 reportedly becomes nuclear excluded [11]. Further, this same lab showed that the phosphomimetic S15D mutation produced a nuclear-excluded Hxk2. Since S15D also gives rise to monomeric Hxk2, they argued that monomeric Hxk2 might be nuclear excluded [11], an idea that has been contested in the literature [25].

Given the uncertainty of the relationship between Hxk2 monomer-dimer regulation and nuclear propensity, we assessed *in vivo* Hxk2 multimerization, as well as the multimerization of the related Hxk1 and Glk1. We performed three co-purification assays from yeast cells expressing two differentially tagged forms of either Hxk1, Hxk2, or Glk1. In all three cases, the V5-purified versions were able to co-purify the GFP-tagged Hxk2, providing evidence for *in vivo* interaction (*i.e.,* the formation of a dimer or higher-order multimer) (Fig 1E). We further found evidence for heterodimer or multimer formation between Hxk1 and Hxk2 (S1G Fig), which also co-purify in high-content studies from yeast [19]. Given that all three hexokinases can multimerize, yet Hxk1 and Glk1 do not undergo a nuclear translocation, this alone is unlikely to explain the differences in their nuclear propensities.

### Dynamics of Hxk2 nuclear partitioning in response to glucose starvation

We further examined Hxk2 nuclear partitioning in response to glucose starvation by performing a time-course image analysis of Hxk2-GFP. In response to low glucose, we observed significantly increased nuclear Hxk2-GFP within 15 minutes, with nuclear accumulation further increasing at 4 and 8 hours before declining at the 24-hour mark (Fig 2A-C; S2A-C Fig). In prolonged starvation conditions, nuclear accumulation of Hxk2-GFP began to diminish, likely due to an increase in protein turnover/degradation as evidenced by the accumulation of free-GFP breakdown products that begins at 8 h post starvation and increases after 24 h of starvation (S2A-D Fig).

**Figure 2.**
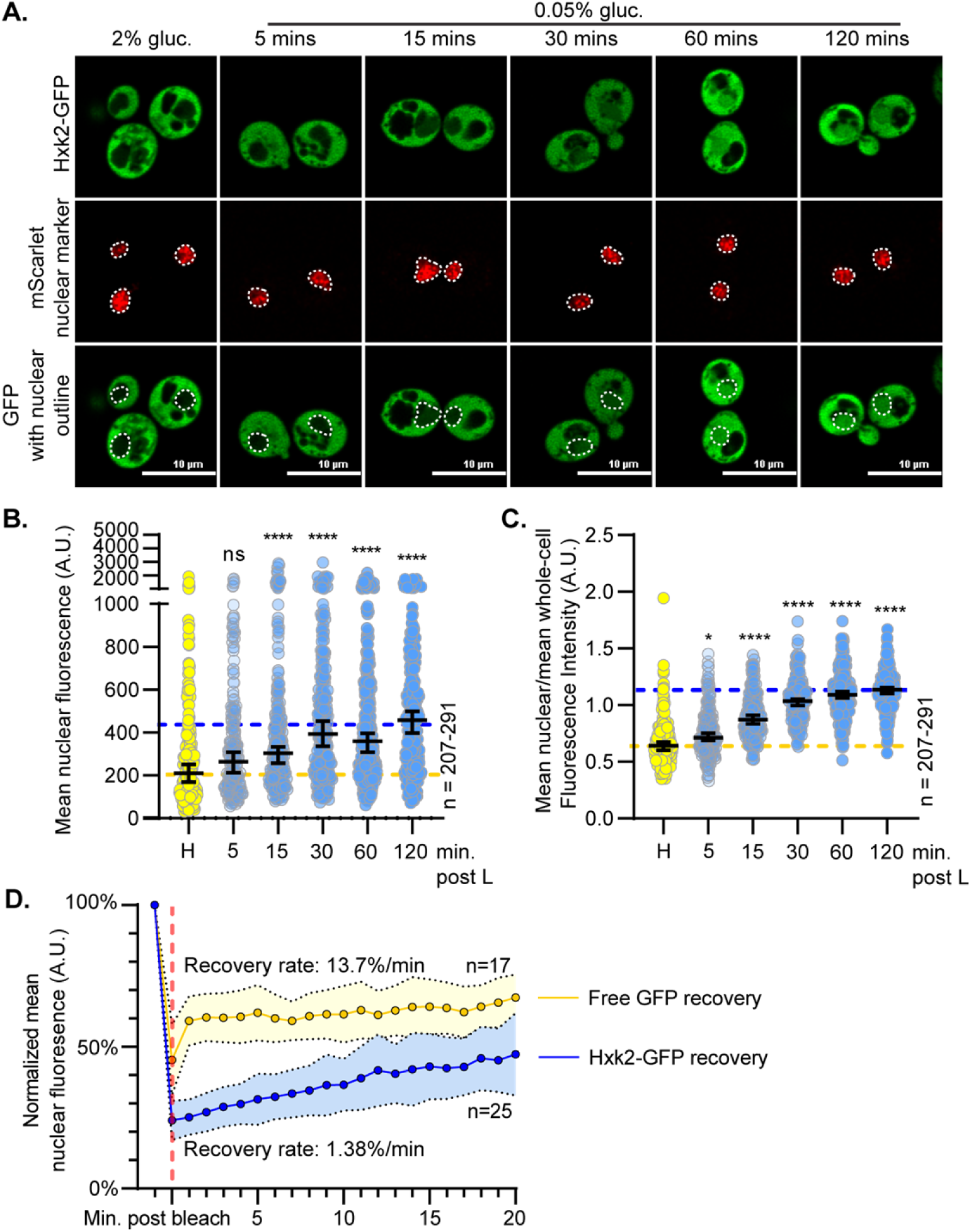
Hxk2 increases its nuclear propensity upon shift to glucose starvation conditions. (A) Confocal microscopy of GFP-tagged Hxk2 expressed from a CEN plasmid under the control of its own promoter in *hxk2*Δ cells. Co-localization with the nucleus is determined based on overlap with the Tpa1-mScarlet nuclear marker, and a dashed white line indicates the nucleus. (B-C) Automated quantification of images shown in panel A to measure (B) mean nuclear fluorescence or (C) the ratio of the mean nuclear/whole-cell fluorescence. Horizontal black lines show the median, and error bars indicate the 95% confidence interval. Dashed yellow and blue lines represent the median value for Hxk2-GFP in high and low glucose, respectively. Kruskal-Wallis statistical analysis with Dunn’s post hoc test was performed to determine if the values obtained post low-glucose shift were statistically different from those obtained in high-glucose conditions. (D) Quantification of FRAP experiments done with cells expressing GFP-tagged Hxk2 from CEN plasmids under the control of its own promoter, and free GFP from CEN plasmids under the control of the *TEF1* promoter. The post-bleaching recovery rate was calculated by measuring the slope of the linear portion of each graph. Blue and gold dots represent the percentage of nuclear fluorescence recovered for Hxk2-GFP and free GFP, respectively. The vertical, red dashed line represents the time point at which nuclear ROI bleaching occurred.

To determine the exchange rate between nuclear and cytosolic Hxk2, we performed fluorescence recovery after photobleaching (FRAP) experiments. We bleached the nuclear Hxk2-GFP in glucose-starved cells and monitored recovery of the nuclear signal. In the 20 min following bleaching, we observed only modest recovery of nuclear signal (∼15% of its initial Hxk2-GFP nuclear signal), suggesting a slow regulated exchange between nuclear and cytosolic Hxk2 (Fig 2D, S2E Fig, and S1 Movie). In fact, nuclear Hxk2-GFP recovered only 1.4% of its fluorescence per minute (Fig 2D).

As a control, we monitored GFP nuclear recovery in cells expressing free-GFP. Free-GFP readily transitions between the nucleus and cytosol [40] and recovered its nuclear fluorescence rapidly, recovering ∼14% of its fluorescence in a minute and plateauing at 60% of its original nuclear signal (Fig 2D and S2E Fig). Thus, unlike GFP, which freely diffuses into the nucleus, Hxk2-GFP has a slow, regulated nuclear translocation.

### Modification of Hxk2 at serine 15 does not alter nuclear partitioning in response to glucose starvation, but does prevent dimerization

Hxk2 phosphorylation at S15 is key for the dimer-to-monomer transition [32–34]. Note that this site is sometimes referred to as S14 in the literature because proteolytic processing removes the Hxk2 N-terminal methionine [41]. Others have reported that in glucose-starvation conditions, phosphorylation at S15 results in nuclear-excluded Hxk2 [11]. However, using confocal microscopy and quantification, we found that Hxk2 with S15 mutated to alanine or the phospho-mimetic aspartic acid (Hxk2^S15A^ and Hxk2^S15D^, respectively) had equivalent nuclear fluorescence to wild-type Hxk2 in glucose-replete and -starvation conditions (Fig 3A-B). Further, the ability of Hxk2^S15A^ and Hxk2^S15D^ to translocate to the nucleus was regulated in the same way as wild-type Hxk2, occurring predominantly upon glucose restriction (Fig 3A-C). These results suggest that the nuclear propensity of Hxk2 does not depend on S15, contradicting earlier studies [11].

**Figure 3.**
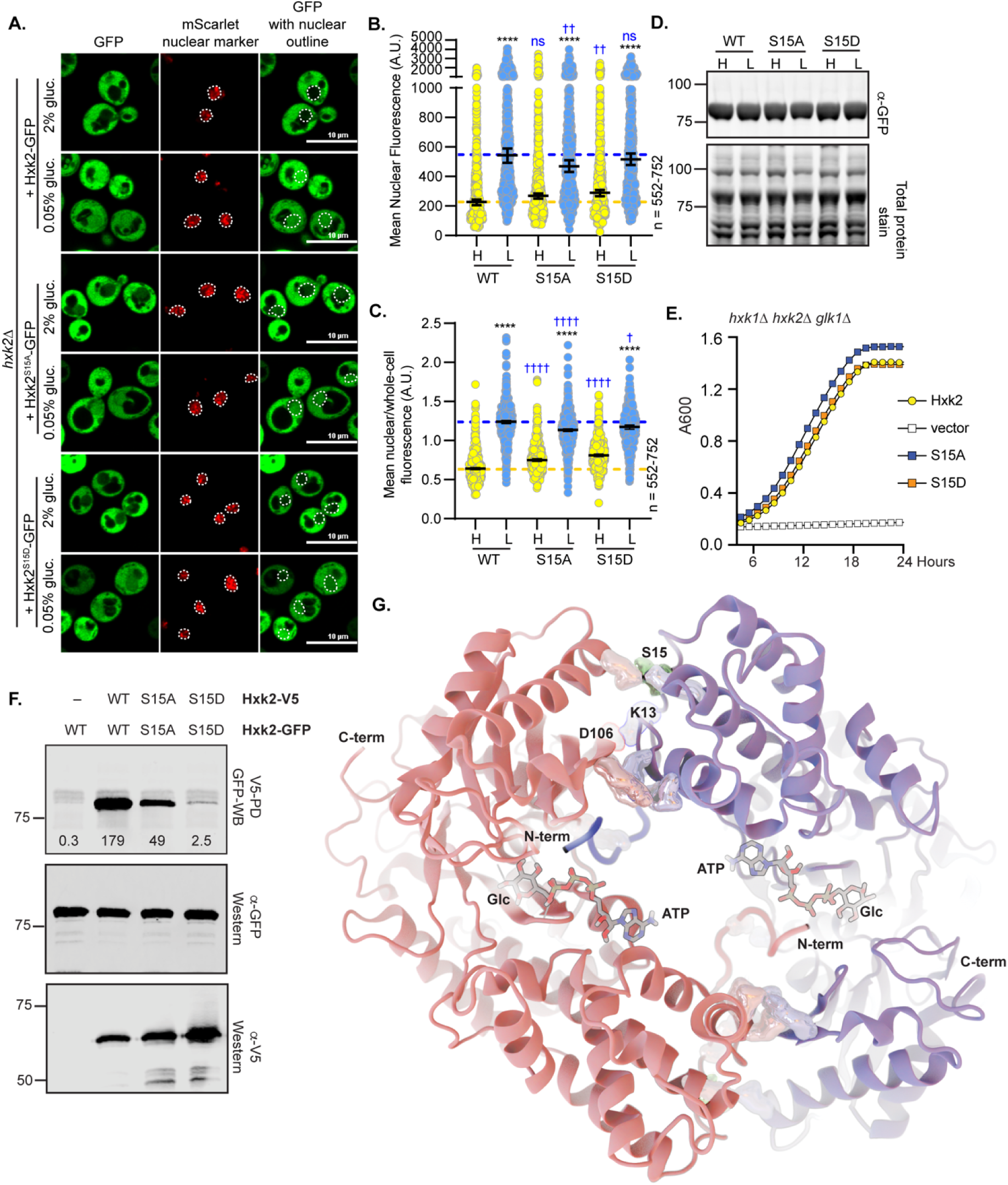
Mutation of Hxk2 at S15 does not alter the regulation of its nuclear translocation but does change its ability to form multimers. (A) Confocal microscopy of GFP-tagged Hxk2 or mutant forms, expressed from a CEN plasmid under the control of the *HXK2* promoter in *hxk2*Δ cells. Co-localization with the nucleus is determined based on overlap with the Tpa1-mScarlet nuclear marker, and a dashed white line indicates the nucleus. (B-C) Automated quantification of the images shown in panel A to measure (B) mean nuclear fluorescence or (C) the ratio of the mean nuclear/whole-cell fluorescence. Horizontal black lines show the median, and error bars indicate the 95% confidence interval. Dashed yellow and blue lines represent the median value for Hxk2-GFP in high and low glucose, respectively. Black asterisks represent statistical comparisons between low and high glucose for a specific *HXK2* allele, and blue daggers represent statistical comparisons between mutant alleles and the corresponding WT Hxk2 in the same medium condition. (D) Immunoblot analyses of Hxk2-GFP from whole cell protein extracts made from cells grown in high glucose or shifted to low glucose for 2 hours. REVERT total protein stain serves as a loading control. (E) Cells lacking all three hexokinase genes (*hxk1Δ hxk2Δ glk1Δ)* were transformed with empty vector or plasmids expressing wild-type Hxk2, Hxk2^S15A^, or Hxk2^S15D^. Cell growth (A_600_) in media containing glucose as the carbon source was monitored for 24 hours. (F) Extracts were prepared from yeast cells expressing the Hxk2 tagged with either V5 or GFP. Hxk2 proteins contained wild-type (WT) S15 or the S15A or S15D mutations. Protein expression was monitored by immunoblotting (bottom two panels). The association of the tagged proteins was assessed by co-immunoprecipitation using anti-V5 beads followed by western blotting with anti-GFP (top panel). Quantitation of the signal in the top panel is shown. (G) The Hxk2-dimer model, with the two monomers shown as pink and blue ribbons, respectively. The glucose molecule and ATP molecules are shown as sticks. The N- and C-terminal tails are marked with “N-term” and “C-term,” respectively. All residues predicted to participate in intermonomer electrostatic interactions are shown as pink and blue metallic surfaces per the associated monomer (see Table 1 for residue numbers). Residue S15 is shown as a metallic green surface.

**Table 1.** Summary of residues at the Hxk2 dimer interface per the homology model of the *Sc*Hxk2 dimer.

| Hxk2 Monomer 1 | Hxk2 Monomer 2 | Interaction type |
| --- | --- | --- |
| VAL 2 | ASP 417 | Salt bridge |
| LYS 7 | GLU 457 | Salt bridge |
| GLN 10 | PHE 105 | Hydrogen bond |
| GLN 10 | THR 107 | Hydrogen bond |
| LYS 13 | ASP 106 | Salt bridge |
| GLN 109 | GLU 356 | Hydrogen bond |
| LYS 111 | ASN 357 | Hydrogen bond |
| LYS 111 | GLU 359 | Salt bridge |
| LYS 111 | ASP 360 | Salt bridge |
| ARG 113 | GLU 359 | Salt bridge |
| ARG 113 | ASP 363 | Salt bridge |
| GLU 141 | LYS 379 | Salt bridge |

We considered the possibility that the role of S15 in Hxk2 localization could be strain specific. Our studies used BY4741-derived yeast, which are related to the S288C genetic background [42], but earlier studies used W303-derived yeast [9–11,36]. We repeated our experiments in W303-derived cells and observed similar Hxk2 nuclear regulation; Hxk2, Hxk2^S15A^, and Hxk2^S15D^ nuclear propensity increased in W303 cells upon glucose starvation, and there was no discernible difference in the nuclear accumulation of these two mutants relative to wild-type Hxk2 (S3A-B Fig). Together, these data demonstrate that S15 does not dramatically alter Hxk2 nuclear regulation.

Hxk2-S15 mutant protein abundances and ability to support growth on glucose in the absence of any other hexokinase were indistinguishable from wild-type Hxk2, demonstrating that they encode stably expressed and catalytically active hexokinases (Fig 3D-E). Importantly, when we assessed *in vivo* multimerization using co-purification from yeast extracts, we found that Hxk2^S15D^ failed to multimerize while Hxk2^S15A^ reduced multimerization, suggesting that each of these mutants diminished Hxk2 dimer formation (Fig 3F). Hxk2^S15D^ strongly disrupts Hxk2 dimerization *in vivo and in vitro*, while Hxk2^S15A^ has a more modest impact *in vitro* [32–35]. Consistent with our observations (Fig 3E), both Hxk2^S15D^ and Hxk2^S15A^ have nearly identical catalytic activities *in vitro* [32–34]. These data confirm that S15 is critical for Hxk2 multimerization, but changes to S15 did not dramatically alter Hxk2 nuclear propensity in wild-type cells. Nuclear translocation was still predominantly driven by glucose starvation.

### Hxk2 homology modeling provides insight into the molecular mechanism of dimerization

To better understand the molecular mechanisms that govern Hxk2 dimerization, we generated a homology model of dimeric *S. cerevisiae* Hxk2 (referred to hereafter as *Sc*Hxk2) based on a crystal structure of the *K. lactis* Hxk1 dimer (PDB 3O1W; referred to hereafter as *Kl*Hxk1) [43]. The 3O1W [43] structure covers almost all the *Kl*Hxk1 sequence without any gaps, including the two N-terminal tails. Crystal structures of many proteins lack N-terminal tails, which are often disordered. In the 3O1W structure, each N-terminus extends into the enzymatic cleft of the opposite *Kl*Hxk1 monomer, which may lock it into a stable position that can be crystallographically resolved. Our *Sc*Hxk2 homology model is similarly complete, including the cleft-bound N-terminal tails (Fig 3G).

To assess which molecular interactions might be responsible for dimerization, we used BINANA [44,45] to identify inter-chain interactions present in the modeled dimer (Table 1 and highlighted in Fig 3G). This analysis identified two primary regions. The first is the N-terminal tail itself. Several charged tail residues participate in salt bridges with the opposite monomer (V2-D417*, K7-E457*, K13-D106*, where an asterisk denotes a residue belonging to the opposite monomer). The Q10 sidechain also forms hydrogen bonds with T107* and F105*. The second region of inter-chain interactions is at the interface between the two distal lobes, where the two monomers also meet. Here K111 forms salt bridges with two residues (E359* and D360*), as does R113 (E359* and D363*). E141 forms a single salt bridge with K379*, and the Q109 and K111 backbones form hydrogen bonds with E356* and N357*, respectively.

Based on our analyses of the *Sc*Hxk2 dimer model, it is not surprising that the two S15 mutants impact dimerization as S15 is near two inter-chain salt bridges (E141-K379* and K13-D106*) (Table 1, Fig 3G). S15 phosphorylation (−2 *e* charge) would change the regional electrostatics, possibly disrupting these dimer-promoting interactions. In contrast, mutation of this site to Ala would preclude phosphorylation and so would preserve the interactions, leaving the dimer intact (Table 1, Fig 3G).

The cleft-bound N-terminal tails also appear to play an important role in promoting *Sc*Hxk2 dimerization (Fig 3G and 4A). Given the crystallographic positions of bound glucose and ATP observed in other structures (e.g., 6PDT [38]), there do not appear to be substantial steric clashes between these bound substrates and the N-terminal tail, even if all three were to occupy the same cleft. We posit that the bound tail might instead be incompatible with catalytic-cleft closure, as is required for glucose phosphorylation. Indeed, our homology model and published crystal structures [43] suggest that hexokinase dimerization generally—and perhaps N-terminal-tail binding specifically—maintains the catalytic cleft in an open conformation. In high glucose concentrations, bound glucose might disrupt N-terminal-tail binding within the catalytic pocket to destabilize the dimer (Fig 4A). Alternatively, in low glucose concentrations, N-terminal-tail binding may prevent glucose binding, encouraging dimerization.

**Figure 4.**
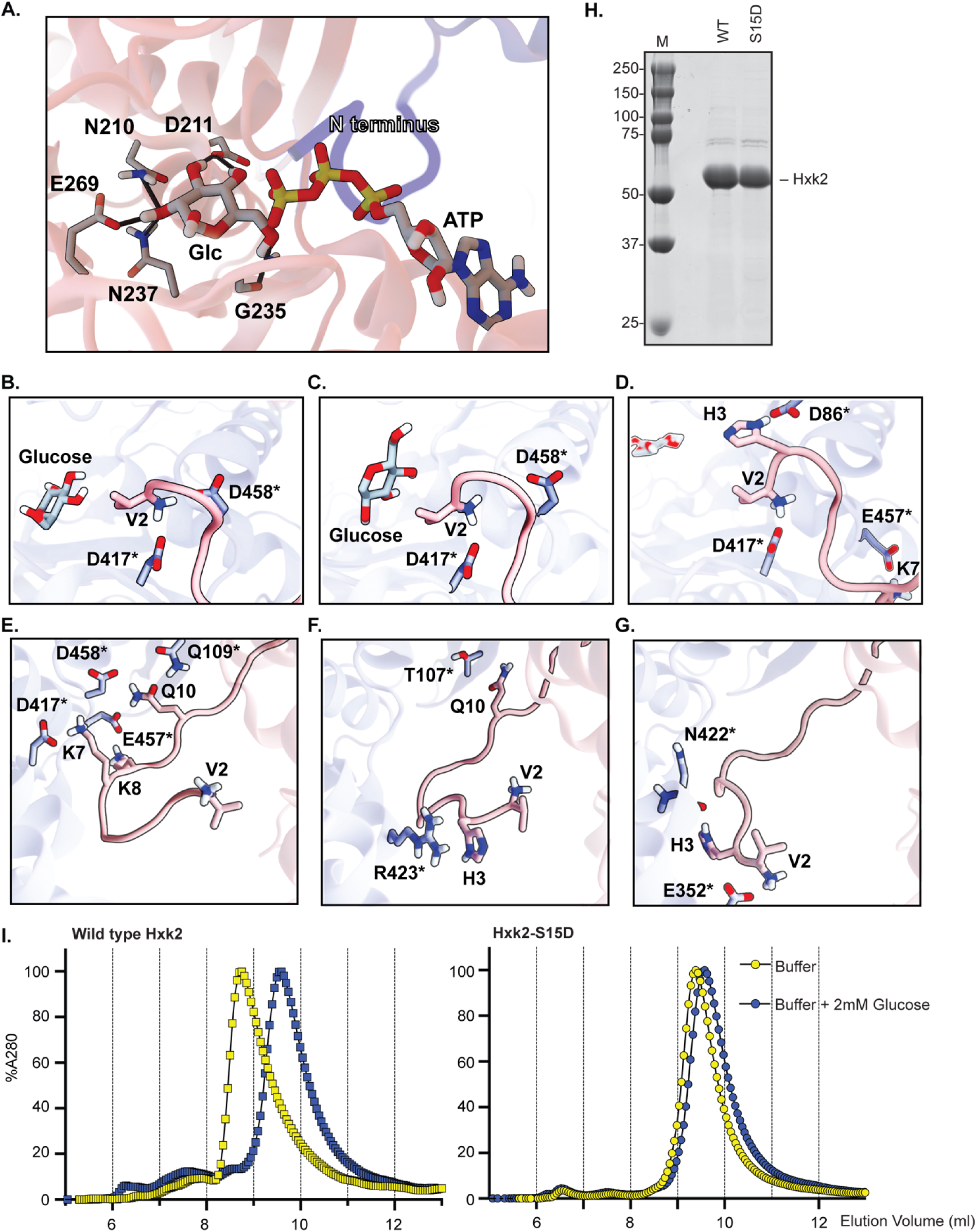
Glucose binding prevents Hxk2 dimer formation. (A) A close-up view of the enzymatic clefts of the Hxk2 dimer model (shown in Fig 3F). The N-terminus from one monomer (in blue) binds in the catalytic pocket of the opposing monomer (in pink). Positioned glucose and ATP molecules are shown in sticks representation. Hydrogen bonds are shown as solid black lines. (B-G) Molecular dynamics simulations demonstrating a possible mechanism of N-terminal-tail dissociation from the enzymatic pocket upon glucose binding. (B) Early in the simulation, the positively charged V2 terminal amine slid between D417* and D458*, forming salt bridges with both. (C) The bound glucose molecule briefly formed hydrophobic contacts with the V2 side chain. (D) Glucose moved to a different location within the enzymatic cleft, and the V2-D458* salt bridge broke. At roughly the same time, a hydrogen bond formed between H3 and D86*. (E) The tail beings to dissociate from the opposite-monomer cleft. K7, which previously formed a salt bridge with E457*, now forms a salt bridge with D417* instead. K8, which did not previously interact with the opposite monomer, forms a salt bridge with E457*. (F) Tail dissociation is stabilized by a cation-π interaction between H3 and R423*, and Q10 shifts to form a hydrogen bond with T107*. (G) Finally, the V2 terminal amine forms a salt bridge with E352*, and H3 forms a hydrogen bond with the backbone carbonyl oxygen of N422*. (H) Yeast Hxk2 proteins (wild type and Hxk2-S15D) were expressed in and purified from *E. coli.* Each protein (5 μg) was resolved by SDS gel electrophoresis and stained with Coomassie blue. (I) Purified, recombinant Hxk2 proteins (WT, square symbols; Hxk2-S15D, round symbols) were resolved by size exclusion chromatography using a buffer with (blue) and without (yellow) glucose.

### Molecular dynamics simulations suggest glucose binding promotes monomerization

To explore this hypothesized glucose/N-terminal-tail antagonism, we performed molecular dynamics simulations of three systems based on our homology model: (1) the *holo* (glucose-bound) *Sc*Hxk2 monomer, (2) the *apo* (ligand-free) *Sc*Hxk2 dimer, and (3) the *holo* (glucose-bound) *Sc*Hxk2 dimer. We did not include ATP in these simulations because we hoped to capture the impact of the initial glucose-binding event before ATP binding and glucose phosphorylation. Simulations of the *holo* (glucose-bound) monomer served to rule out the possibility that starting the simulations in an open conformation alone would allow for glucose dissociation, even when the cleft is not occupied by an N-terminal tail. We performed three independent simulations of the *holo* monomer starting from the open (dimer-like) conformation (1,043 ns total). At no point did the glucose molecule deviate substantially from its initial binding pose (mean RMSD: 2.24 Å; standard deviation: 0.89 Å).

Simulations of the *apo* (glucose-absent) dimer model served to rule out the possibility that the N-terminal tail is prone to dissociate on its own independent of bound glucose, perhaps due to inaccuracies in our homology model. We performed three simulations of the *apo Sc*Hxk2 dimer (1,078 ns total). At no point did any N-terminal tail dissociate from its opposite-monomer cleft.

In contrast, simulations of the *holo* (glucose-bound) *Sc*Hxk2 dimer captured notable glucose and N-terminal-tail dynamics. We performed three simulations of this system. In the first (262 ns total), both N-terminal tails remained associated with their respective opposite-monomer enzymatic clefts. The pose of one glucose molecule also remained stable (mean RMSD: 1.04 Å; standard deviation: 0.25 Å), but the other glucose was highly mobile within the cleft (mean RMSD: 5.11 Å; standard deviation: 2.56 Å). The second simulation of the *holo Sc*Hxk2 dimer (261 ns total) was similar. Both N-terminal tails remained associated with their respective clefts, one glucose remained stable (mean RMSD: 2.45 Å; standard deviation: 0.87 Å), and one glucose was mobile within its cleft (mean RMSD: 5.93 Å; standard deviation: 3.25 Å). Given that we saw little glucose mobility in 1,043 ns of *holo Sc*Hxk2 monomer simulation, we hypothesize that dimerization—and perhaps the opposite-monomer N-terminal tail specifically—discourages stable glucose binding at the enzymatic site.

The third simulation of *holo Sc*Hxk2 dimer (1,000 ns total) explored dimer dynamics on slightly longer timescales. During this simulation, one of the N-terminal tails remained associated with its respective opposite-monomer enzymatic cleft. The glucose molecule in the same cleft was again highly mobile, so much so that it dissociated entirely from the protein. In contrast, the other N-terminal tail dissociated from its enzymatic cleft. The same-cleft glucose molecule was also highly mobile, though it remained in the cleft (average RMSD: 9.60 Å; standard deviation: 2.52 Å). Given that we saw little N-terminal-tail mobility in 1,078 ns of *apo* Hxk2 dimer simulation, we hypothesize that bound glucose encourages N-terminal-tail dissociation.

### Proposed mechanism of initial dimer dissociation

Because the third simulation captured the only observed tail dissociation, we explored its mechanistic details in greater depth. Early in the simulation, the positively charged terminal amine group of V2 is sandwiched between D458* and D417*. Other nearby inter-monomer interactions include the K7-E457* salt bridge and the Q10-T107*, Q10-D106*, and Q10-F105* hydrogen bonds (Fig 4B).

A dissociation cascade begins when the bound glucose molecule transiently reorients to form hydrophobic contacts with the V2 sidechain (Fig 4C). This brief interaction has two consequences. First, it expels the glucose molecule to a cleft-bound position distant from the N-terminal tail. Second, it weakens the V2-D458* interaction, such that V2 interacts more exclusively with D417*. This change permits a new hydrogen bond to form between H3 and D86*, possibly partially compensating for the weakened V2-D458* interaction (Fig 4D).

However, the V2-D417* and H3-D86* interactions break after ∼130 ns, causing the N-terminal tail to move away from the cleft. This large-scale movement impacts other inter-monomer interactions as well. Within about 100 ns, the K7-E457* interaction breaks, permitting the formation of a new K7-D417* interaction. Q10 also prefers Q109* and D458* as hydrogen-bond partners rather than T107*, D106*, and F105* as before. Amino acid K8, which did not previously interact with the opposite monomer, forms a strained electrostatic interaction with E457* (Fig 4E).

Two subsequent conformational rearrangements further destabilize the N-terminal-tail/cleft association. The first rearrangement occurs after another ∼230 ns. H3, now free of D86*, forms a cation-π interaction with R423*, and Q10 returns to its hydrogen bond with T107* (Fig 4F). The second rearrangement occurs after another ∼270 ns when V2 forms a salt bridge with E352* that further distances it from the cleft. The H3-D86* cation-π interaction breaks to accommodate this change, and H3 instead forms a hydrogen bond with the backbone carbonyl oxygen atom of N422* (Fig 4G).

These dynamics support the hypothesis that an N-terminal tail and glucose molecule do not simultaneously occupy the same enzymatic cleft. Of note, neither of the two *Kl*Hxk1-dimer crystal structures that fully resolve the N-terminal tail (PDB IDs 3O1W and 3O4W [43]) include a bound glucose molecule. The one dimer structure that does include a glucose molecule (PDB ID 3O5B) does not have a fully resolved N-terminal tail, perhaps because the tail cannot bind (and be stabilized by) the opposite-monomer enzymatic cleft.

### Chromatography confirms that glucose binding to Hxk2 promotes monomer formation

Our simulations captured only one N-terminal-tail dissociation event, so we cannot guarantee that this dissociation pathway is the most common. We further note that the simulation did not run long enough to capture the full dissociation of the entire dimer, which may occur on far longer (perhaps computationally intractable) timescales.

To verify that glucose alone promotes dimer dissociation independently of S15 phosphorylation, we used chromatography to biochemically examine the multimerization of purified recombinant Hxk2 and Hxk2^S15D^ in the absence or presence of glucose. Using nickel affinity columns, we obtained highly purified Hxk2 and Hxk2^S15D^ from *E. coli* (Fig 4H). As a control we first confirmed that Hxk2^S15D^ promotes monomerization in the absence of glucose. In ion exchange chromatography experiments, wild-type Hxk2 eluted as a single peak with an ∼8.8 mL elution volume. Hxk2^S15D^ took longer to elute from the column (elution volume of ∼9.5 mL), consistent with a smaller size than wild-type Hxk2 (Fig 4I, compare yellow curves). This change in elution profile is consistent with wild-type Hxk2 forming a larger, dimeric complex than the Hxk2^S15D^ mutant in the absence of glucose. In contrast, Hxk2^S15D^ cannot dimerize unless much higher enzyme concentrations are used [33] and so represents monomeric Hxk2 [32,33].

To assess the impact of glucose, we preincubated Hxk2 with glucose in the absence of ATP, locking the enzyme in a glucose-bound state, before performing ion exchange chromatography. In the presence of glucose, Hxk2 migrated more slowly and eluted as a single peak at ∼9.5 mL, the same elution profile observed with the monomeric Hxk2^S15D^ (Fig 4I, compare blue curve for WT Hxk2 to yellow Hxk2^S15D^ curve). Unlike wild-type Hxk2, the elution profile of the monomeric Hxk2^S15D^ mutant did not change upon adding glucose, as expected since it was already monomeric (Fig 4I, compare yellow and blue curves for Hxk2^S15D^). Since these assays were done without ATP it allowed unphosphorylated glucose to remain in the binding pocket. They are thus directly comparable to the simulations, which similarly omitted ATP.

These results confirm that glucose binding disrupts dimer formation. They support earlier biochemical studies demonstrating that glucose binding (1) promotes Hxk2 monomer formation and (2) greatly decreases the dimerization association constant. The Hxk2 dimer had a K_a_ of 1.2 x 10^7^/M when glucose was absent versus 4.5 x 10^4^/M in the presence of glucose. A similar mutant to the one used here, Hxk2^S15E^, had a dimer K_a_ of 1.3 x 10^4^/M without glucose present, nearly 1000-fold lower than wild-type Hxk2 in the same conditions [33]. We propose a model that incorporates our MD simulations and biochemical findings with the earlier work from the Kriegel lab: (1) Glucose and N-terminal tail binding in the Hxk2 catalytic cleft are mutually incompatible. (2) N-terminal tail docking into the catalytic cleft of the opposing monomer helps stabilize Hxk2 dimerization. (3) Modification of the N-terminal S15 disrupts the dimer-interface by disrupting N-tail-catalytic pocket association between opposing monomers.

### The Hxk2 N-terminal tail, thought initially to contain an NLS, is required for Hxk2 dimer formation and nuclear exclusion

To further assess the role of the N-terminal tail, we made a mutant Hxk2 that lacked amino acids 7-16 (referred to as Hxk2^Δ7-16^). This same mutation has been used in the past and referred to as the “<u>w</u>ithout regulatory function” (WRF) mutation based on its reported inability to regulate glucose-repression of the *SUC2* gene [46]. Studies from the same lab suggest that Hxk2^Δ7-16^ retains full catalytic function but fails to bind Mig1 and does not localize to the nucleus, suggesting that a subset of these amino acids constitute an Hxk2 nuclear localization sequence [9,46].

However, our earlier work demonstrates that the Hxk2^Δ7-16^ mutant retains partial catalytic activity (∼50% that of WT Hxk2) and maintains glucose repression of the *SUC2* gene [4], refuting the earlier claim that this mutant fails to repress gene expression [9,46]. We were surprised to find that Hxk2^Δ7-16^ localized to the nucleus in both glucose-replete and glucose-starvation conditions, with significantly higher mean nuclear fluorescence than wild-type Hxk2 and a dramatic increase in the mean nuclear to whole-cell fluorescence ratios (Fig 5A-C). Thus, rather than serving as a nuclear localization sequence, the Hxk2 N-terminus appears critical for maintaining its glucose-regulated, nuclear exclusion.

**Figure 5.**
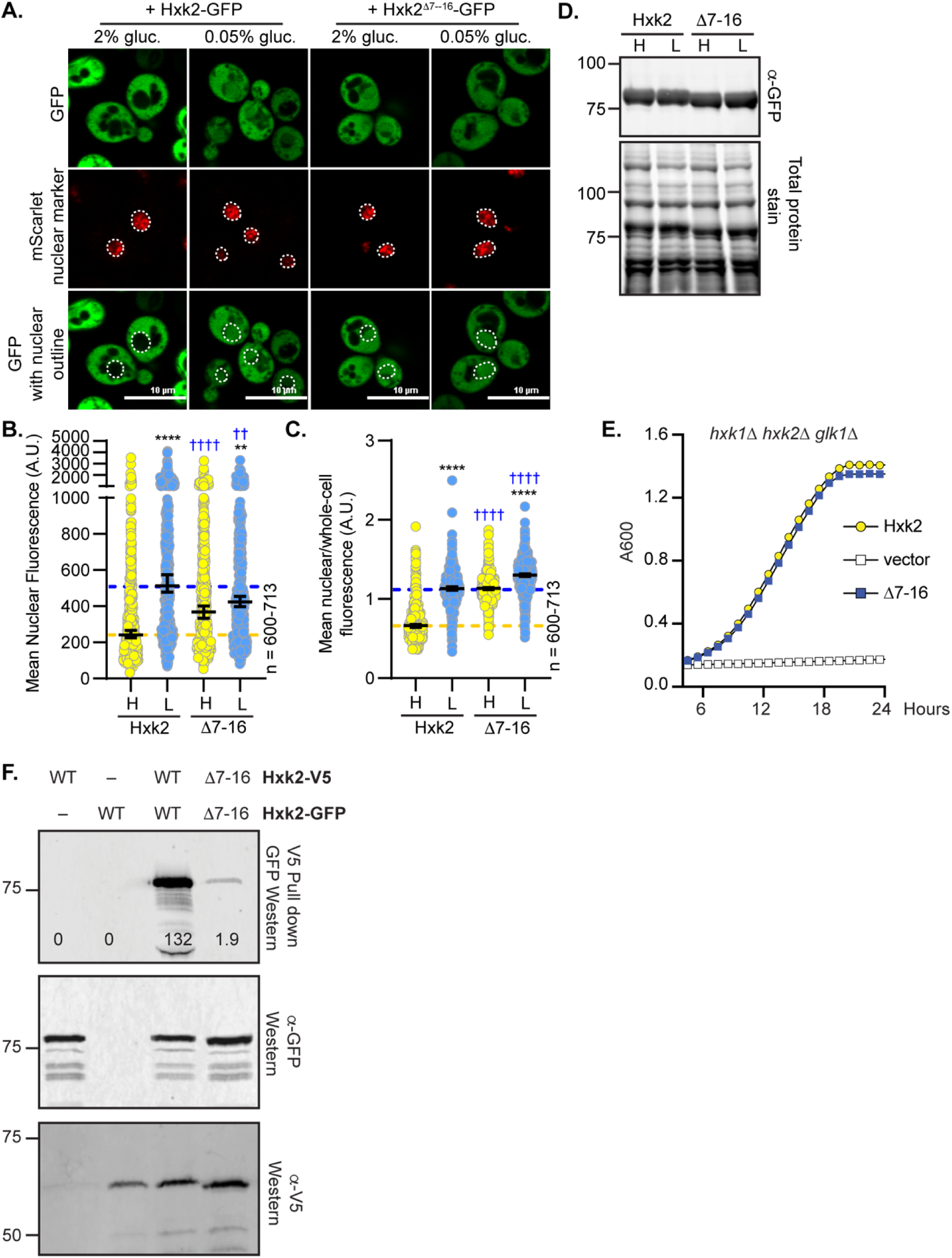
Deleting the N-terminal amino acids 7-16 results in constitutive Hxk2 nuclear localization, prevents Hxk2 dimerization but maintains catalytic function. (A) Confocal microscopy of GFP-tagged Hxk2 or Hxk2^Δ7-16^ expressed from a CEN plasmid under the control of the *HXK2* promoter in *hxk2*Δ cells. Co-localization with the nucleus is determined based on overlap with the Tpa1-mScarlet nuclear marker, and a dashed white line indicates the nucleus. (B-C) Automated quantification of the images shown in panel A to measure (B) mean nuclear fluorescence or (C) the ratio of the mean nuclear/whole-cell fluorescence. Horizontal black lines show the median, and error bars indicate the 95% confidence interval. Dashed yellow and blue lines represent the median value for Hxk2-GFP in high and low glucose, respectively. Black asterisks represent statistical comparisons between low and high glucose for a specific *HXK2* allele, and blue daggers represent statistical comparisons between mutant alleles and the corresponding WT Hxk2 in the same medium condition. (D) Immunoblot analyses of Hxk2-GFP in whole-cell protein extracts made from cells grown in high glucose or shifted to low glucose for 2 hours. REVERT total protein stain of the membrane serves as a loading control. (E) Cells lacking all three hexokinase genes (*hxk1Δ hxk2Δ glk1Δ)* were transformed with plasmid vector or plasmids expressing wild-type Hxk2 or Hxk2-Δ7-16, as indicated. Cell growth (A_600_) in media containing glucose as the carbon source was monitored for 24 hours. (F) To assess multimerization, we prepared extracts from yeast cells expressing the untagged Hxk2 or Hxk2 tagged with either V5 or GFP. Hxk2 proteins contained either the wild-type (WT) N-terminus or the Δ7-16 deletion. Protein expression was monitored by western blotting (bottom two panels). The association of the tagged proteins was assessed by co-immunoprecipitation using anti-V5 beads followed by western blotting with anti-GFP (top panel).

Hxk2 proteins lacking these ten N-terminal amino acids were still stable and functional, as confirmed by immunoblotting and their ability to support robust growth on glucose for cells lacking endogenous hexokinase (Fig 5D-E). However, the Hxk2^Δ7-16^ mutant did not copurify a differentially tagged version of Hxk2^Δ7-16^, suggesting it cannot multimerize (Fig 5F). This result is not surprising considering our modeling and simulations, which suggest the Hxk2 N-terminal tail is necessary for dimer formation (Fig 3G and Table 1).

There are three lysines among amino acids 7-16, and lysines are often critical for NLS function [47]. To assess the specific roles of these lysines, we made site-directed mutants at lysines 7, 8, and 13, converting these residues to alanine (referred to as Hxk2^K7,8,13A^). This triple mutant recapitulated the constitutive nuclear localization observed for Hxk2^Δ7-16^ (S4A-C Fig). Rather than forming an NLS, this region helps maintain glucose-induced nuclear exclusion of Hxk2.

### K13 in the Hxk2 N-terminal tail is required for Hxk2 dimer formation and glucose regulation of nuclear localization

We found that the Hxk2^Δ7-16^ and Hxk2^K7,8,13A^ mutants prevented dimerization and circumvented glucose-regulated nuclear translocation, giving rise to a pool of constitutively nuclear Hxk2. However, disrupting dimerization is not sufficient to drive constitutive nuclear localization of Hxk2 because Hxk2^S15D^, which prevents dimerization, largely retains glucose-regulated nuclear exclusion (Fig. 3A-G and S3A-B Fig). Interestingly, amino acids 7-16 in Hxk2 are perfectly conserved in Hxk1, but Hxk1 does not accumulate in the nucleus in response to glucose starvation (Fig 6A and Fig 1A-D), and analogous mutations in Hxk1 to those under study for Hxk2 did not result in accumulation of nuclear Hxk1 (S5A-C Fig).

**Figure 6.**
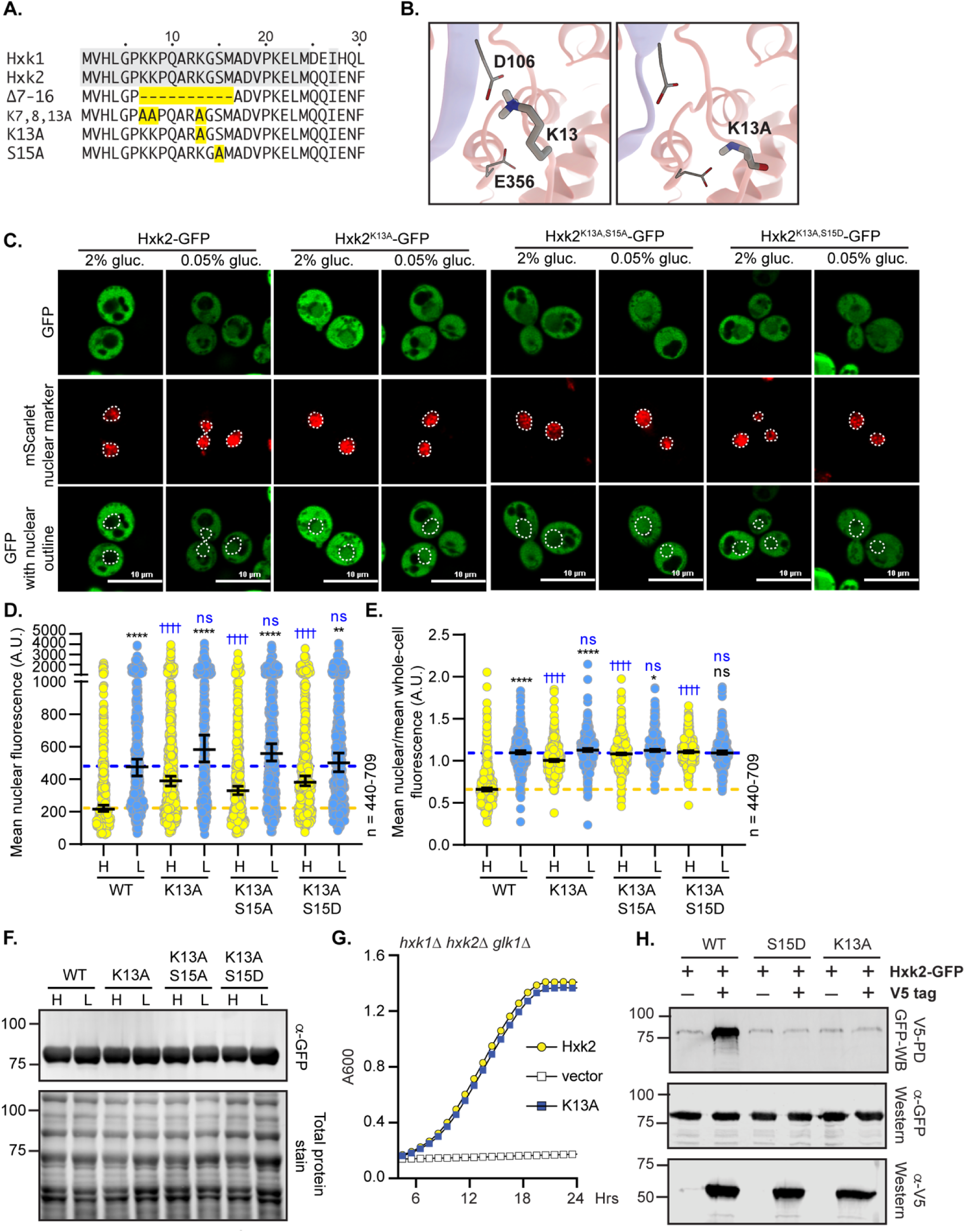
Mutation of K13 to alanine in the Hxk2 N-terminal tail results in constitutive Hxk2 nuclear localization and prevents Hxk2 dimerization but maintains catalytic function. (A) Sequence alignment of the Hxk2 N-terminal tail with Hxk1, highlighting key mutated residues. The first 30 residues of Hxk1 and Hxk2 are shown with identical residues shaded. Select mutations in this region are shown in yellow. (B) Modeled K13 interactions. The two Hxk2 monomers are shown as pink and blue ribbons. In the first panel, a close-up view of K13 interactions with E365 (same monomer) and D106 (opposing monomer). In the second panel, an alanine substitution at K13 cannot form the same electrostatic interactions. (C) Confocal microscopy of GFP-tagged Hxk2 or the mutant alleles expressed from a CEN plasmid under the control of the *HXK2* promoter in *hxk2*Δ cells. Co-localization with the nucleus is determined based on overlap with the Tpa1-mScarlet nuclear marker, and a dashed white line indicates the nucleus. (D-E) Automated quantification of imaging shown in (C) to measure (D) mean nuclear fluorescence or (E) the ratio of the mean nuclear/whole-cell fluorescence. Horizontal black lines show the median, and error bars indicate the 95% confidence interval. Dashed yellow and blue lines represent the median value for Hxk2-GFP in high and low glucose, respectively. Black asterisks represent statistical comparisons between low and high glucose for a specific *HXK2* allele, and blue daggers represent statistical comparisons between mutant alleles and the corresponding WT Hxk2 in the same medium condition. (F) Immunoblot analyses of Hxk2-GFP in whole-cell protein extracts made from cells grown in high glucose or shifted to low glucose for 2 hours. (G) Cells lacking all three hexokinase genes (*hxk1Δ hxk2Δ glk1Δ)* were transformed with plasmid vector or plasmids expressing wild-type Hxk2 or Hxk2^K13A^, as indicated. Cell growth (A_600_) in media containing glucose as the carbon source was monitored for 24 hours. (H) Extracts were prepared from yeast cells expressing Hxk2-GFP and Hxk2 with or without the V5 as indicated. Hxk2 proteins contained either the wild-type (WT) sequence or the S15D or K13A mutations. Protein expression was monitored by western blotting (bottom two panels). The association of the tagged proteins was assessed by co-immunoprecipitation using anti-V5 beads followed by western blotting with anti-GFP (top panel). Quantitation of the signal in the top panel is shown.

To refine the region responsible for glucose-regulated nuclear localization, we considered our *Sc*Hxk2 dimer homology model, which suggests the N-terminal-tail residues K7 and K13 form important salt bridge interactions that may stabilize the dimer (Figs 3F and 6B). Of these two sites, K13 is likely dimethylated or sumoylated in Hxk2 [48,49] (Fig 6B) but ubiquitinated in Hxk1 [50]; this possible differential regulation of Hxk1 and Hxk2 could account for their distinct nuclear localization patterns. We assessed the localization of GFP-tagged Hxk2^K13A^, a single-point mutation likely to disrupt the modeled K13-D106 electrostatic interaction observed in our *Sc*Hxk2 homology model (Fig 6B, center panel). Like the more severe Hxk2^Δ7-16^ and Hxk2^K7,8,13A^ mutants, Hxk2^K13A^ was nuclear localized in both glucose-replete and glucose-starved conditions (Fig 6C-E). It had elevated nuclear fluorescence in 2% glucose-grown cells and an elevated nuclear to whole-cell fluorescence ratio that was indistinguishable from that of wild-type Hxk2 under glucose-starvation conditions. Further, there was little change in the mean nuclear to whole-cell fluorescence ratio of Hxk2^K13A^ between glucose-grown or -starved cells (Fig 6E), demonstrating that this mutant bypasses glucose-inhibition of nuclear localization. Mutation of Hxk1 to generate the analogous K13A mutant did nothing to change the distribution of Hxk1, which remained cytosolic in all glucose growth conditions (S5A-C Fig).

As with the other related alleles tested, Hxk2^K13A^ encodes a stable and functional protein, has catalytic activity, and permits robust growth on glucose when expressed as the only hexokinase in cells (Fig 6F-G). As we suspected, the K13A mutation disrupted multimer formation as effectively as Hxk2^S15D^ (Fig 3G and 6H); the V5-tagged Hxk2^K13A^ from yeast extracts could not copurify GFP-tagged Hxk2^K13A^ (Fig 6H).

It should be noted that the K13A mutation and dimer disruption allow for constitutive access to the S15 phosphorylation site, which others have suggested might control nuclear translocation [11]. To determine if S15 phosphorylation influenced the misregulated localization of the K13A mutant, we combined K13A with S15A or S15D, which prevent phosphorylation or mimic phosphorylation, respectively. Neither of these new double mutants – Hxk2^K13A,S15A^ nor Hxk2^K13A,S15D^ – showed any difference in nuclear localization compared to the Hxk2^K13A^ mutation alone (Fig 6C-E). Thus, we propose that S15 phosphorylation is dispensable for nuclear translocation and/or retention of Hxk2^K13A^.

We next mutated D106, the residue that pairs with K13 to form a salt bridge at the Hxk2 dimer interface (Fig 3G). Interestingly, the Hxk2^D106A^ mutant, which would presumably break the interaction with K13 and allow it to be accessible for modification and/or binding, maintained normal Hxk2 cytosol-nuclear partitioning in both glucose-replete and -starvation conditions (Fig 7A-C). However, as anticipated, the D106A mutant was unable to form multimers *in vivo* (Fig 7D), supporting the idea that K13 should be accessible in this mutant. These data demonstrate that generating the Hxk2 monomer is not sufficient to drive nuclear localization of Hxk2. In addition, access to K13, as would be expected to occur in any mutant that breaks the dimer, is not enough to stimulate nuclear localization, supporting a model where post-translational modification or some aspect specific to the lysine at this position is required for glucose regulation of Hxk2’s nuclear propensity.

**Figure 7.**
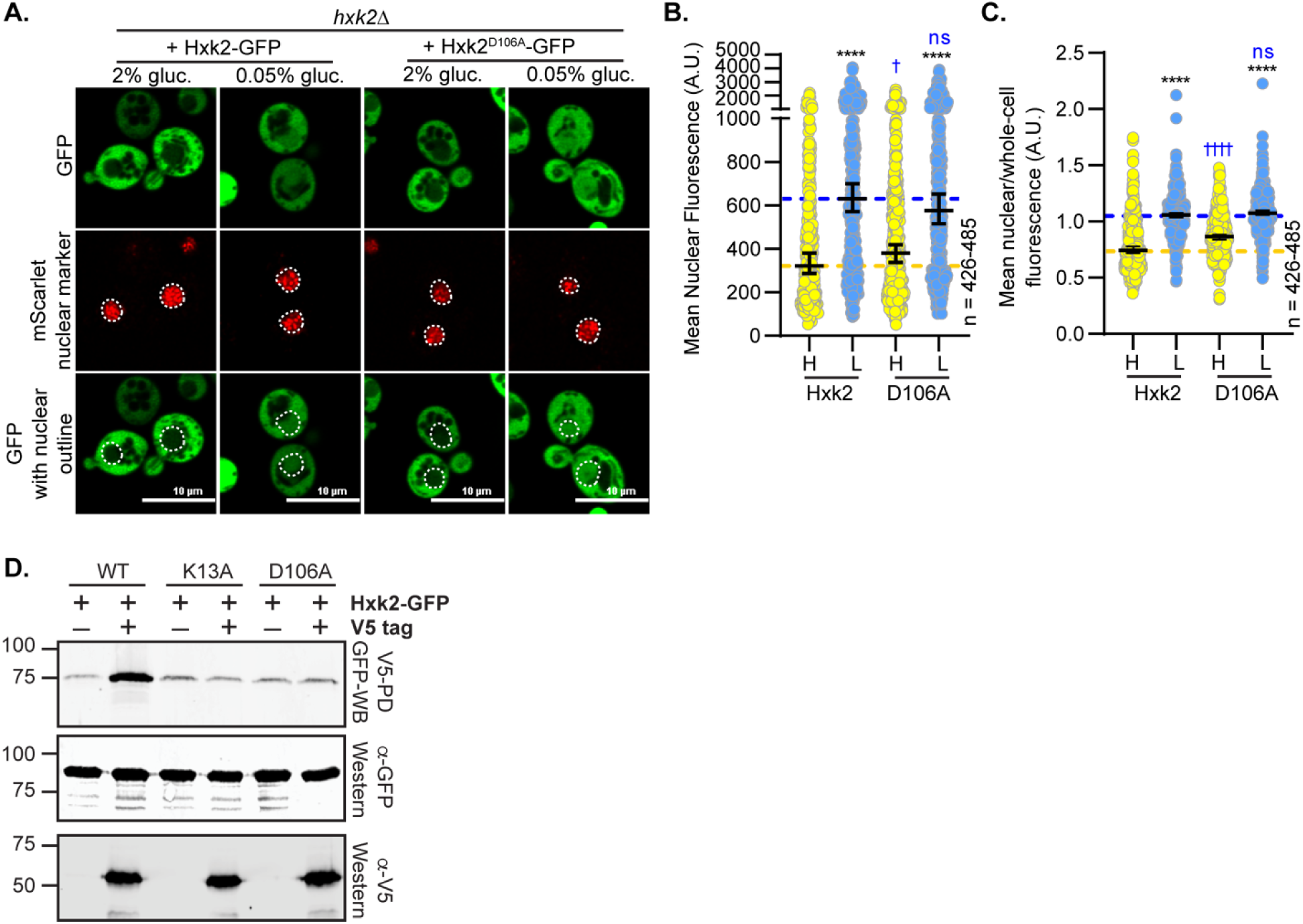
Mutation of D106, which interacts with K13, does not alter Hxk2 nuclear localization but does prevent Hxk2 dimerization. (A) Confocal microscopy of GFP-tagged Hxk2 or the mutant alleles expressed from a CEN plasmid under the control of the *HXK2* promoter in *hxk2*Δ cells. Co-localization with the nucleus is determined based on overlap with the Tpa1-mScarlet nuclear marker, and a dashed white line indicates the nucleus. (B-C) Automated quantification of imaging shown in (A) to measure (B) mean nuclear fluorescence or (C) the ratio of the mean nuclear/whole-cell fluorescence. Horizontal black lines show the median, and error bars indicate the 95% confidence interval. Dashed yellow and blue lines represent the median value for Hxk2-GFP in high and low glucose, respectively. Black asterisks represent statistical comparisons between low and high glucose for a specific *HXK2* allele, and blue daggers represent statistical comparisons between mutant alleles and the corresponding WT Hxk2 in the same medium condition. (D) Extracts were prepared from yeast cells expressing Hxk2-GFP and Hxk2 with or without the V5 as indicated. Hxk2 proteins contained either the wild-type (WT) sequence or the K13A or D106A mutations. Protein expression was monitored by western blotting (bottom two panels). The association of the tagged proteins was assessed by co-immunoprecipitation using anti-V5 beads followed by western blotting with anti-GFP (top panel). Quantitation of the signal in the top panel is shown.

### Tda1, but not Snf1 or Mig1, is required for Hxk2 nuclear accumulation

Many studies have identified Hxk2 S15 as a phosphorylated residue [31,51–56], lending strong support to the idea that this residue is regulated. This site is also conserved perfectly in Hxk1, where it is also phosphorylated [51–57]. Multiple kinases are linked to Hxk2 S15 phosphorylation, including PKA, Snf1, and Tda1 [11,31,35]. In addition to being important for Hxk2 phosphorylation, the Snf1 kinase, its substrate Mig1, and Reg1 (an activator of the PP1 protein phosphatase Glc7 that controls Snf1 activity) are thought to interact with Hxk2 to facilitate its nuclear translocation, where they form a large complex that controls Hxk2’s alleged moonlighting function as a transcriptional regulator [7,9,11,58,59].

We examined the impact of Snf1, Mig1, and the upstream Reg1 on Hxk2 nuclear propensity. In the absence of Snf1 or Mig1, there was little change in Hxk2 nuclear localization in glucose-replete or -starvation conditions (Fig 8A-C). Upon glucose starvation, *snf1*Δ and *mig1*Δ cells had slightly reduced or elevated mean nuclear fluorescence, respectively, compared to WT cells, but these changes were either not significant or just past the significance threshold (Fig 8A-B). When the ratio of nuclear to whole-cell fluorescence was considered, *mig1*Δ cells were not different than WT under any condition, while *snf1*Δ cells had increased Hxk2 nuclear balance in high glucose conditions (Fig 8C). These results counter earlier findings, which suggested Mig1 is required for Hxk2 nuclear translocation [9].

**Figure 8.**
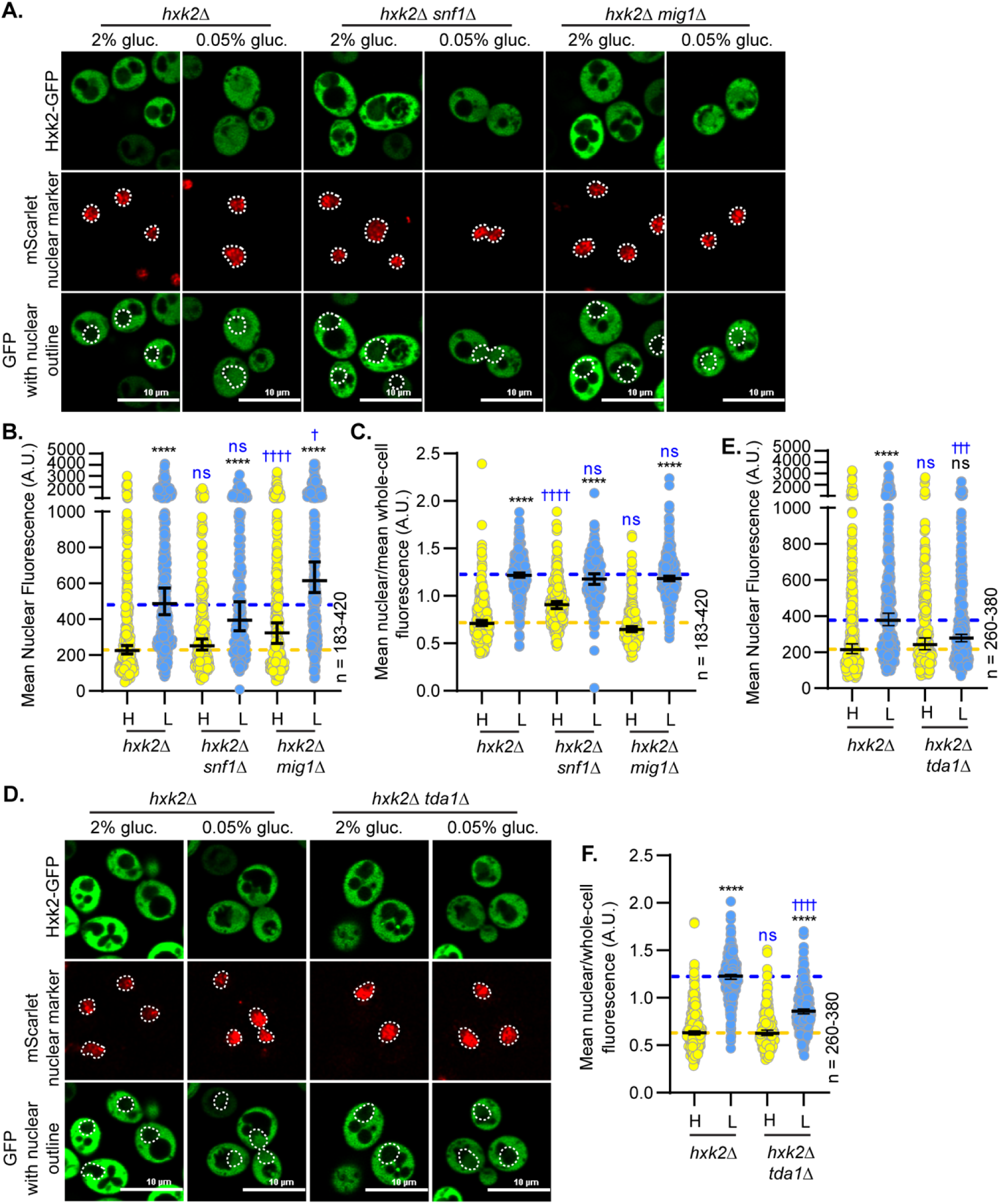
Hxk2 nuclear localization is not regulated by Snf1 or Mig1 but is controlled by the Tda1 kinase. (A and D) Confocal microscopy of GFP-tagged Hxk2 expressed from a CEN plasmid under the control of the *HXK2* promoter in cells lacking *HXK2* alone or further missing (A) *SNF1* or *MIG1* or (D) *TDA1*. Co-localization with the nucleus is determined based on overlap with the Tpa1-mScarlet nuclear marker, and a dashed white line indicates the nucleus. (B-C; E-F) Automated quantification of the images shown in panels A or D to measure (B or E, respectively) mean nuclear fluorescence or (C or F, respectively) the ratio of the mean nuclear/whole-cell fluorescence. Horizontal black lines show the median, and error bars indicate the 95% confidence interval. Dashed yellow and blue lines represent the median value for Hxk2-GFP in high and low glucose, respectively. Black asterisks represent statistical comparisons between low and high glucose for a specific *HXK2* allele, and blue daggers represent statistical comparisons between mutant alleles and the corresponding WT Hxk2 in the same medium condition.

We attempted to recapitulate the reported co-purification of Hxk2 with Mig1. However, we could not capture any HA-tagged Mig1 above the weak background signal of bead binding using a Hxk2-V5 pulldown (S6A Fig). Although negative data, they are consistent with our other observations suggesting *mig1*Δ does not impact Hxk2 nuclear translocation.

Surprisingly, the loss of Reg1 completely abolished Hxk2 nuclear translocation in response to glucose starvation (S6B-D Fig). Thus, Reg1 — and likely by extension the PP1 phosphatase, Glc7 — are required for Hxk2 nuclear accumulation in response to glucose starvation. We will examine the impact of this regulator on Hxk2 in future studies because we have not yet determined if Reg1 directly influences Hxk2 phosphorylation or indirectly regulates Snf1 or other Glc7-Reg1 substrates.

Elegant studies from Kriegel and Kettner demonstrate that the Tda1 kinase is important for phosphorylating Hxk2 S15, thereby preventing Hxk2 dimerization [25,60,61]. Consistent with a role for Tda1 in Hxk2 regulation, glucose-starvation-induced Hxk2 nuclear accumulation was severely dampened in *tda1*Δ cells (Fig 8D-F). Recent biochemical fractionation studies found that Tda1 becomes nuclear localized in glucose-starved cells [62]. In our microscopy experiments, mNeonGreen-tagged Tda1 signal was very low in glucose-replete conditions with a 3-fold increase in the mean nuclear and whole-cell fluorescence upon glucose starvation (Fig 9A-C). However, the nuclear-to-whole-cell fluorescence ratio was only modestly different in glucose-grown versus -starved cells demonstrating that the nuclear-cytosolic balance of Tda1 is maintained in these two conditions (Fig 9D). Immunoblot analyses also showed a significant increase in Tda1-mNG abundance in response to glucose starvation (Fig 9E).

**Figure 9.**
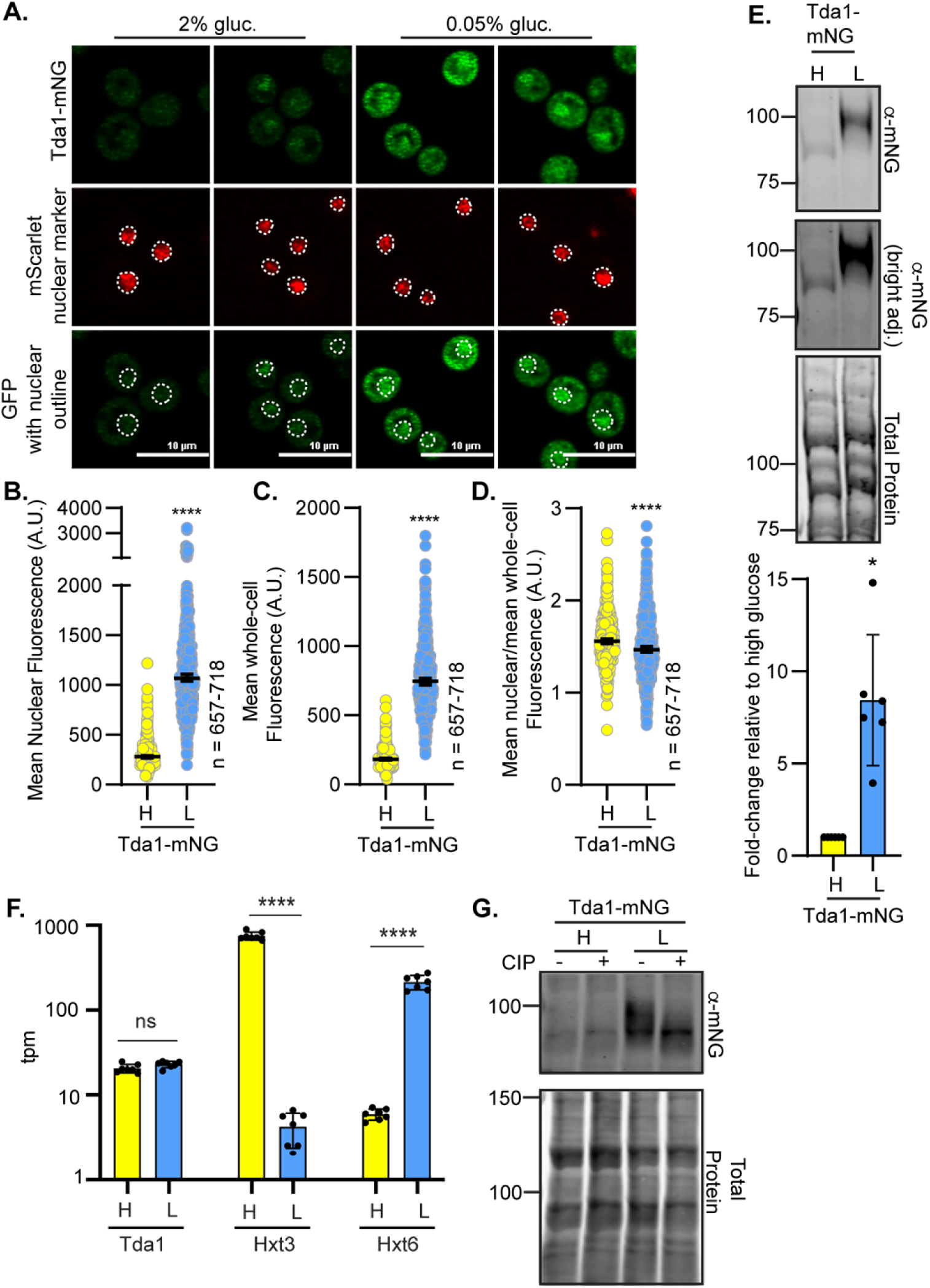
Tda1 protein abundance and phosphorylation are increased in response to glucose starvation. (A) Confocal microscopy of chromosomally integrated Tda1-mNG. Co-localization with the nucleus is determined based on overlap with the Tpa1-mScarlet nuclear marker, and a dashed white line indicates the nucleus. (B-D) Automated quantification of the images shown in panel A to measure (B) mean nuclear fluorescence, (C) mean whole-cell fluorescence, or (D) the ratio of the mean nuclear/whole-cell fluorescence. Horizontal black lines show the median, and error bars indicate the 95% confidence interval. A Student’s t-test was used to assess significance (****= p-value < 0.0001). (E) Immunoblot analyses of Tda1-mNG in whole-cell protein extracts made from cells grown in high glucose or shifted to low glucose for 2 hours. REVERT total protein stain of the membrane serves as a loading control. One representative blot from 6 biological replicates is shown. Quantification of Tda1-mNG abundance is presented as a fold-change in Tda1 levels after a 2 h shift to low glucose in comparison to glucose-grown cells (high glucose). (F) Yeast mRNA abundance (transcripts per million mapped reads; tpm) was measured by RNAseq in cells grown in high glucose (H) or two hours after shifting to low glucose, L. The mean tpm values (±SD) of multiple replicates for each condition are plotted for three genes: *TDA1*, *HXT3,* and *HXT6.* Statistical differences between the values in high and low glucose are indicated. (G) Whole-cell extracts were made from strains expressing Tda1-mNG and grown in either 2% glucose, H, or shifted for 2 h into low glucose, L (0.05% glucose). Extracts were either treated with calf intestinal alkaline phosphatase (CIP) or incubated in the same conditions without enzyme (mock), resolved by SDS-PAGE, and immunoblotted with anti-mNG antibody. REVERT total protein stain is shown as a loading control. Molecular weights are shown on the left side in kDa.

Increased Tda1 protein levels were not driven by elevated *TDA1* transcription, as RNAseq analyses showed no change in *TDA1* transcript abundance for cells grown in 2% or 0.05% glucose conditions for 2 hours (Fig 9F), unlike the glucose-responsive transcripts of hexose transporters 3 and 6 (*HXT3* and *HXT6*, respectively) shown as controls. As expected [63,64], *HXT3* transcripts were reduced upon glucose starvation, and *HXT6* transcripts increased (Fig 9F). The increased Tda1 protein could be due to altered translation rates or a change in Tda1 protein stability/regulation.

Consistent with altered regulation, our immunoblots showed a sizeable shift to a slower-mobility Tda1 form in low-glucose conditions (Fig 9E). This mobility change was due to phosphorylation, as incubation with phosphatase gave rise to a single band with the same migration pattern as Tda1 in glucose-grown cells (Fig 9G). Based on these data, we propose that Tda1 is required for Hxk2 nuclear accumulation in glucose-starved cells. Tda1 is a glucose-regulated kinase, and glucose starvation increases its protein abundance and phosphorylation. Elevated Tda1 protein levels likely reflect post-transcriptional regulation since Tda1 transcript abundance is unchanged in glucose-starved cells.

We next examined the impact of Hxk2 mutations on Tda1-dependent Hxk2 nuclear translocation. As before, Hxk2 accumulation in the nucleus was severely blunted in glucose-starved *tda1*Δ cells (Fig 8D-F and Fig 10A-C). However, the S15D mutation restored glucose-starvation induced Hxk2 nuclear localization in *tda1*Δ cells (Fig 10A-C). The balance of Hxk2^S15D^ nuclear-to-whole-cell fluorescence ratios in *tda1*Δ cells was very similar to that of Hxk2 in wild-type cells but was shifted to be slightly higher in Hxk2^S15D^ *tda1*Δ cells in high glucose conditions. Unlike Hxk2^S15D^, the Hxk2^K13A^ mutation largely bypassed both Tda1 and glucose-starvation regulation; Hxk2^K13A^ was still nuclear in glucose-grown *tda1*Δ cells, though its overall nuclear abundance was modestly reduced compared to glucose starvation conditions (Fig 10B), and Hxk2^K13A^ retained a much larger nuclear pool than WT Hxk2 in both glucose-grown and -starved conditions (Fig 10A-C).

**Figure 10.**
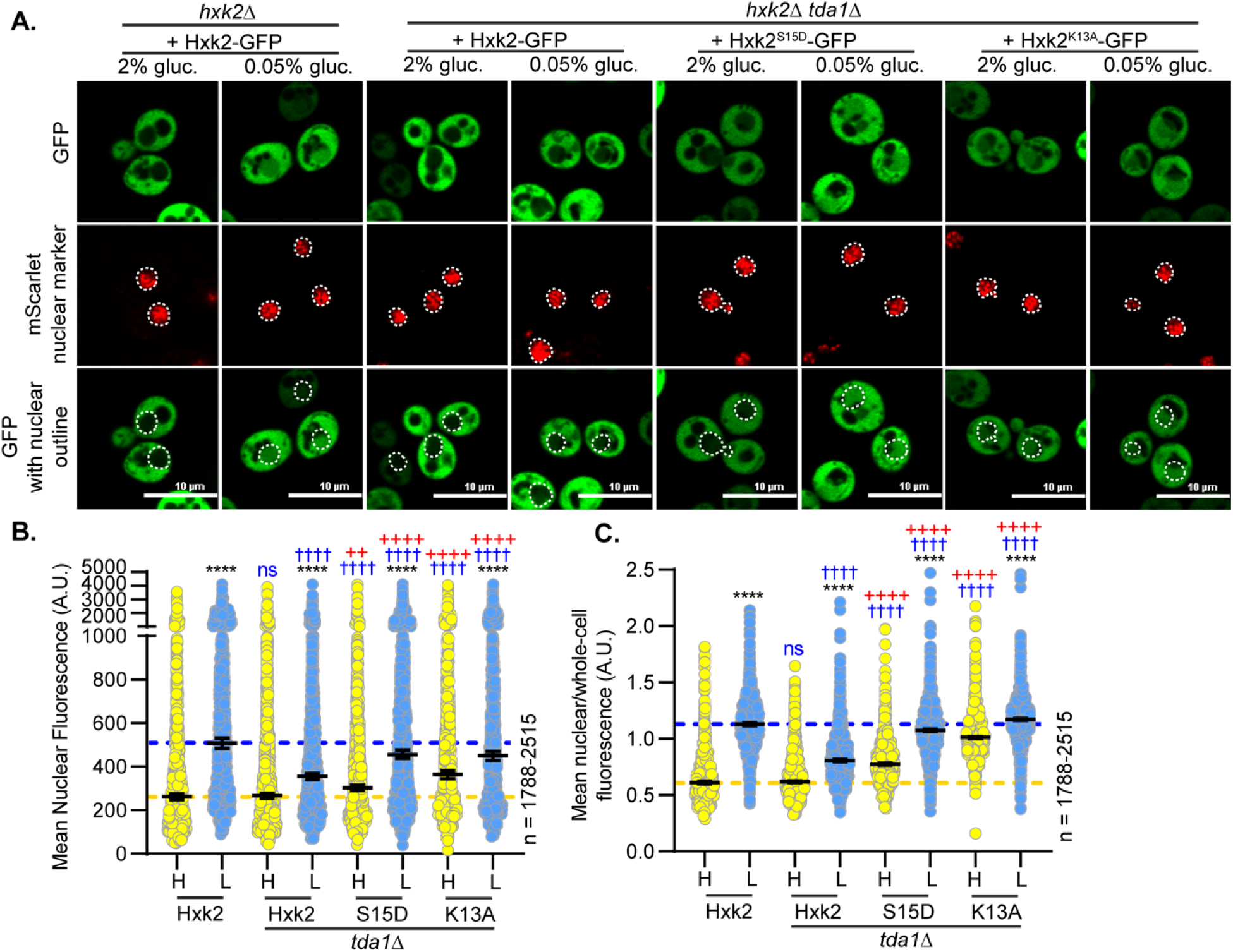
Mutating Hxk2 S15 to aspartic acid rescues the impaired nuclear localization caused by a *tda1*Δ. (A) Confocal microscopy of GFP-tagged Hxk2 and S15D mutant expressed from a CEN plasmid under the control of the *HXK2* promoter in cells lacking *HXK2* alone or with the additional deletion of *TDA1*. Co-localization with the nucleus is determined based on overlap with the Tpa1-mScarlet nuclear marker, and a dashed white line indicates the nucleus. (B-C) Automated quantification of the images shown in panel A to measure (B) mean nuclear fluorescence or (C) the ratio of the mean nuclear/whole-cell fluorescence. Horizontal black lines show the median, and error bars indicate the 95% confidence interval. Dashed yellow and blue lines represent the median value for Hxk2-GFP in high and low glucose, respectively. Black asterisks represent statistical comparisons between low and high glucose for a specific *HXK2* allele. Blue daggers represent statistical comparisons between mutant alleles and the corresponding WT Hxk2 in the same medium condition. Red crosses represent statistical comparisons between WT Hxk2 and Hxk2^S15D^ in the *hxk2*Δ *tda1*Δ background in the same media condition.

Although the Hxk2^S15D^ mutation restored nuclear localization in glucose-starved *tda1*Δ cells, this mutation had a different effect in *snf1*Δ cells (S7A-C Fig). We observed a higher nuclear-to-whole-cell ratio in glucose-grown *snf1*Δ cells expressing Hxk2^S15D^ than those expressing wild-type Hxk2 (S7A and S7C Figs). Interestingly, this increase in nuclear-to-whole-cell fluorescence was not driven by higher mean nuclear fluorescence, which is the same for Hxk2^S15D^ and Hxk2 in *snf1*Δ or wild-type cells (S7B Fig). Together, these findings suggest the S15D phosphomimetic mutation bypasses Tda1 regulation, restoring glucose regulated Hxk2 nuclear accumulation. However, unlike the Hxk2^K13A^ mutant, the Hxk2^S15D^ mutation does not alter Hxk2 localization in glucose-grown cells.

Since Hxk2^S15D^, Hxk2^D106A^, and Hxk2^K13A^ all disrupt Hxk2 dimerization *in vivo*, yet only Hxk2^K13A^ prevents glucose-regulated nuclear localization, Hxk2 dimerization cannot be the only facet controlling Hxk2 nuclear partitioning. Hxk2^S15D^, which favors monomeric Hxk2, is not required for the block to Hxk2 nuclear partitioning in glucose-grown cells; rather, it appears important for glucose-starvation-induced Hxk2 nuclear localization. In contrast, Hxk2^K13A^, which also favors monomeric Hxk2, bypasses glucose-dependent regulation of Hxk2 nuclear partitioning and is no longer reliant on phosphorylation at S15.

### Role of Hxk2 in regulating glucose-repression of gene expression

Earlier studies proposed that Hxk2 is a subunit of a DNA-bound repressor complex and so regulates gene expression and in particular glucose repression [7]. To analyze the role of Hxk2 in gene expression and glucose repression, we performed RNA-seq analyses of the yeast transcriptome in wild type and *hxk2Δ* cells grown in high glucose and after shifting to glucose limiting conditions (0.05% glucose) for two hours. RNA samples were prepared in triplicate and the log2+1 ratios (low glucose/high glucose) for the mean abundance of each mRNA were plotted as a function of mRNA abundance (Fig 11A-D). As reported previously [4,65], the yeast transcriptome undergoes large-scale changes in mRNA abundance in response to glucose limitation with >15% of the transcripts showing a 4-fold or higher change in abundance (Fig 11A). Notably, large changes (both increases and decreases) in the abundance of the hexose transporter (*HXT*) mRNAs were observed as well as a striking decrease in the abundance of the ribosomal protein mRNAs. A comparable pattern, scale and scope of mRNA abundance changes were observed in the RNA samples generated from *hxk2Δ* cells (Fig 11B), demonstrating that the Hxk2 protein is not required for the global transcriptional response to changes in glucose abundance. Replotting these values to show the ratio of mRNA abundance (*hxk2*Δ/wild type) in high glucose (Fig 11C) and low glucose (Fig 11D) further demonstrates that very few mRNAs are impacted by the deletion of *HXK2*. Interestingly, the few genes (*SUC2, HXT1, HXK1*) that have been reported to be affected by *HXK2* [8,63,66] were standouts in high glucose and were not representative of the bulk of glucose regulated genes. A violin plot of the log2+1 ratios from this experiment demonstrated the large, and similar, response to glucose limitation in both wild type and *hxk2Δ* cells (Fig 11E). By comparison, the difference between wild-type and h*xk2*Δ cells under these conditions was modest with nearly no 2-fold or greater changes in transcript abundances (Fig 11E). Finally, we plotted the mean mRNA abundance of the top 20 glucose-repressed genes in high and low glucose in wild type and *hxk2*Δ cells (Fig 11F). Glucose repression and derepression of these mRNAs was not affected by deletion of the *HXK2* gene, even for three genes whose promoters are bound by Mig1 protein [22]. These data demonstrate that Hxk2 is not a transcriptional regulator, nor does it play a role in regulating glucose repression.

**Figure 11.**
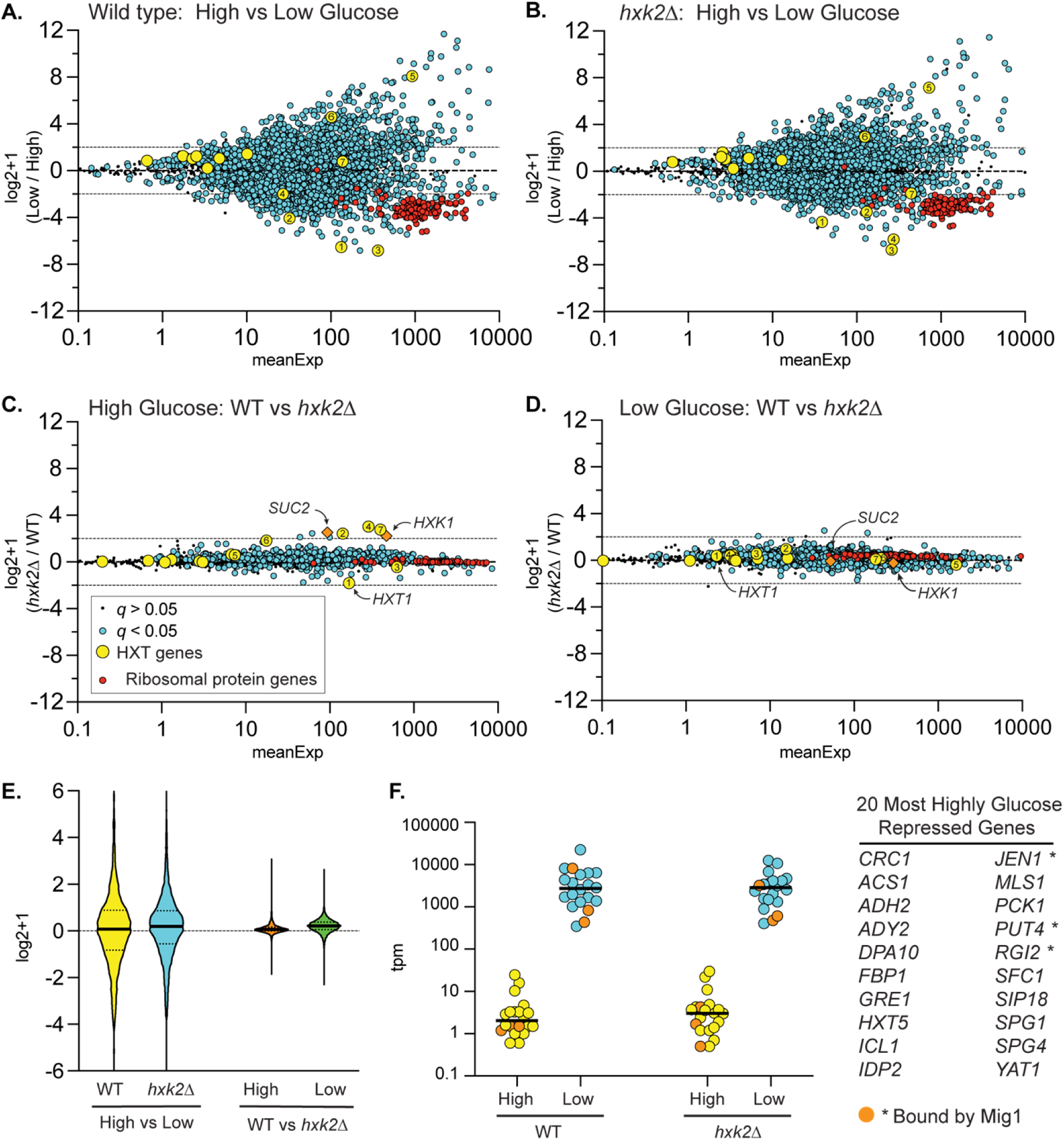
Effect of Hxk2 on the transcriptional response to glucose limitation. Total RNA was collected from 3 independent cultures of wild-type cells or *hxk2Δ* cells grown to mid-log phase in high glucose or two hours after shifting to low glucose. mRNA abundance (tpm) for all genes was determined by RNAseq and is plotted as the mean expression level (x-axis) versus the log2+1 value of the ratio of the mean value in low glucose divided by the mean value in high glucose (A and B) in wild type (A) or *hxk2Δ* cells (B). The same data are replotted, showing the log2+1 ratio of *hxk2* and wild-type cells in high (C) and low glucose (D) conditions. In each plot, statistical significance was defined as a false discovery rate (q) less than 0.05. Log2+1 values deemed significant were plotted as blue circles, while those failing to meet this threshold were plotted as smaller black dots (see key). *HXT* gene mRNAs were plotted as yellow circles with the number referring to the specific *HXT* gene. Ribosomal mRNA genes were plotted as red circles. The known Hxk2-regulated genes, *SUC2* and *HXK1,* were plotted as orange diamonds. (E) Violin plot of these data showing the relative magnitude of glucose limitation (high vs. low) and *hxk2* deletion (WT vs. *hxk2Δ*). (F) The top 20 glucose-repressed genes (listed at right) are plotted based on their TPMs from both high glucose (yellow) and low glucose (blue) conditions. Three of these genes (indicated by * in the table at right and shown as orange-filled circles on the plot) are reported to have Mig1 bound in their promoters based on ChIP-Seq.

## Discussion

Glucokinases and hexokinases are central metabolic regulators that control glucose conversion to G6P, the critical first step in glycolysis. In addition to cytosolic functions in glycolysis, each enzyme of this pathway can accumulate in the nucleus. For mammalian and plant hexokinases, nuclear accumulation typically occurs in glucose starvation or other stress conditions [26,27,67–69]. What is the nuclear function of glycolytic enzymes? Perhaps they act together in nuclear glycolysis to regulate a nuclear pool of ATP or NADH [61,70]. Alternatively, could they have a nuclear role in regulating gene expression or other facets of nuclear biology? The nuclear function of many glycolytic enzymes remains mechanistically unclear [70]. However, for a handful of examples, there is evidence for diverse roles that involve altering transcription factor function, associating with RNA polymerase III, interacting with DNA, regulating nuclear ubiquitin ligases, stimulating DNA polymerase, and protecting telomeres [71–75]. In some instances, the metabolic products generated by glycolytic enzymes may be the active nuclear component rather than the proteins [76]. Much remains to be discovered about the “moonlighting” nuclear activities of glycolytic enzymes.

In studies of hexokinase localization, *S. cerevisiae* and *C. albicans* hexokinase 2 have been outliers. Their nuclear accumulation reportedly occurs in glucose-replete rather than starvation conditions [9–11,36,37]. However, the methodologies used in early imaging studies involved cells pre-incubated in a medium that would induce glucose-starvation. This medium likely confounded the interpretation of these studies, especially if the incubation time in the glucose-starvation medium was not regimented [9–11,36,37].

### Contrasts between our findings and the existing model for Hxk2 nuclear regulation

Here we report high-resolution, live-cell, confocal microscopy of fluorescently tagged Hxk2 performed in high (2% glucose) and low (0.05% glucose) glucose medium. Our findings support a new model for Hxk2 nuclear translocation and suggest that in yeast Hxk2 accumulates in the nucleus under glucose-starvation conditions, not glucose-replete conditions. This is consistent with a model of conserved, glucose-regulated hexokinase nuclear translocation spanning the ∼150 million years of evolution that separate yeasts and humans.

Our data contradict the existing model of regulation for Hxk2 nuclear translocation in other ways as well. In contrast to earlier models, we find that: (1) Mig1 is not required for Hxk2 nuclear translocation; (2) the previously reported NLS/Mig1 binding site (amino acids 7-16) is not required for Hxk2 nuclear translocation but instead helps maintain a glucose-regulated, nuclear-excluded pool of Hxk2; (3) Snf1 is not required for the glucose-regulated nuclear accumulation of Hxk2; (4) Reg1 is important for Hxk2 nuclear translocation, since in the absence of Reg1, Hxk2 fails to become nuclear localized even in glucose starvation conditions; (5) phosphorylation of S15, though a key regulator of the Hxk2 monomer-dimer balance, is not required for the glucose-regulated Hxk2 nuclear accumulation; and (6) except for modest changes in a handful of transcripts, Hxk2 is not required to maintain glucose-repression of transcription.

### A new model for regulation of Hxk2 nuclear translocation

Based on our studies, we propose a new model for glucose-regulated Hxk2 nuclear accumulation (Fig 12). In a glucose-replete environment, the Glc7-Reg1 phosphatase is active, maintaining Snf1 in an inactive state [77,78]. The Tda1 kinase, though transcribed, is in low abundance in cells grown in 2% glucose, suggesting it is either not translated or generates an unstable protein product. Under these conditions, Hxk2 S15 is not phosphorylated, and Hxk2 exists as a balance between monomer and dimer species [32,33].

**Figure 12.**
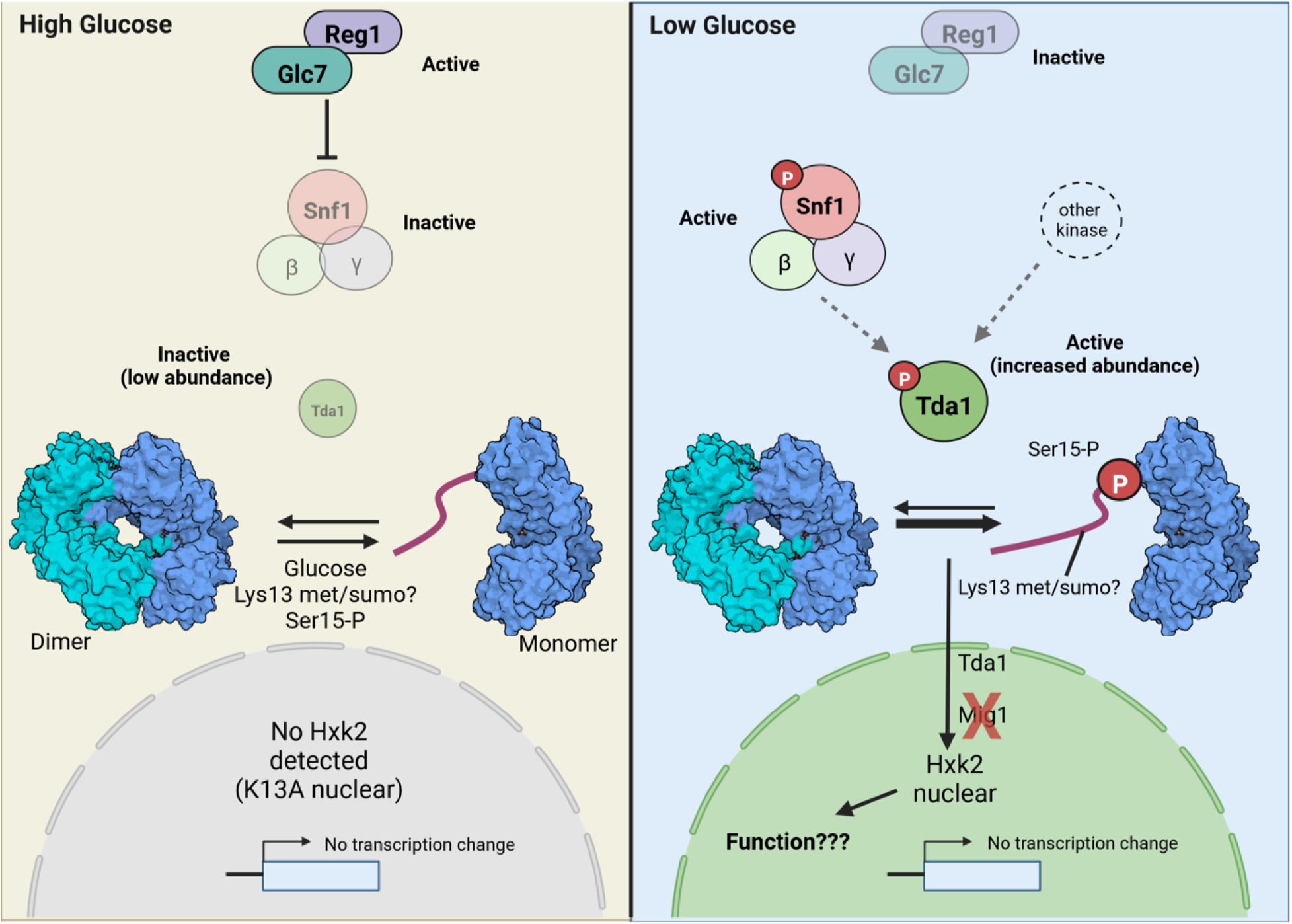
Model for glucose-regulation of Hxk2 nuclear accumulation and dimerization. (A) A schematic of key regulatory factors and their function when cells are grown in glucose replete (referred to as high glucose) conditions. For cells grown in glucose, the Reg1-Glc7 protein phosphatase complex is active, and this dephosphorylates Snf1 to keep it inactive. The impact of this regulation on Tda1 is currently unclear. The Tda1 kinase is present in cells at very low levels and is an inactive kinase. Glucose binding to the Hxk2 dimer (the two Hxk2 monomers are shown in light and dark blue, respectively) stimulates monomer formation, as does mutation of the enzyme at S15 or K13. Under these conditions, no nuclear Hxk2 is detected unless the K13A mutant is present, which somehow inactivates the glucose-induced block to Hxk2 nuclear accumulation. From the RNA-seq analyses, Hxk2 does not regulate glucose-repressed gene expression. (B) A schematic of key regulatory factors and their function when cells are glucose starved (referred to as low glucose). Under these conditions, Reg1-Glc7 is inactivated. However, we find that Reg1 is needed for Hxk2 nuclear translocation in this condition, but the mechanism underlying this requirement is unclear. Snf1 kinase is phosphorylated and activated. It is unclear if Snf1 is responsible for phosphorylating Tda1 to activate it kinase. However, it is tempting to speculate that this could be one mode of Snf1 regulation in this pathway. In glucose starvation conditions, the abundance of Tda1 increases dramatically, and Tda1 is phosphorylated and activated. In these conditions, phosphorylation of Hxk2 by Tda1 at S15 regulates the Hxk2 dimer to monomer transition. Tda1 is required for Hxk2 nuclear translocation, but Mig1 and Snf1 are not needed. Hxk2 phosphorylation at S15 reduces the capability of Hxk2 to dimerize by more than 1000x [32]. This stabilization of the monomeric species could allow for additional modifications at the Hxk2 N-terminus. Notably, K13 is reportedly dimethylated or sumoylated [48,49], and either of these modifications could contribute to regulated Hxk2 nuclear import. Once in the nucleus, Hxk2 does not significantly impact transcription regulation. Based on our RNA-seq analyses, there is little difference between the expression profiles of WT vs. *hxk2*Δ cells glucose starvation conditions. The function of nuclear Hxk2 remains to be determined.

We find that Hxk2 shifts to a monomer when glucose binds, confirming earlier studies that demonstrate a dramatic increase in the association constant of Hxk2 dimers when glucose is present [33]. Molecular dynamics simulations provide insight into why glucose might favor Hxk2 monomer formation. In our simulations, bound glucose impacts multiple electrostatic interactions between the opposite monomer’s N-terminal tail and the catalytic pocket, which may ultimately promote Hxk2 dimer dissociation. Our experimental evidence confirms that the N-terminal tail is critical for dimerization. Deleting amino acids 7-16 or mutating K13 to alanine both give rise to monomeric Hxk2. Disrupting the K13-D106 salt bridge at the dimer interface by alanine substitution at D106 also breaks the dimer but does not stimulate nuclear accumulation of Hxk2 in glucose replete conditions.

Hxk2 monomer-dimer balance is likely an important facet of Hxk2 regulation, but it is not responsible for Hxk2 nuclear translocation. Not all mutations that induce monomer formation stimulate Hxk2 nuclear accumulation. For example, Hxk2^S15D^ and Hxk2^D106A^ both fail to dimerize but retain glucose-regulated nuclear translocation. On the other hand, Hxk2^K13A^ and an Hxk2 mutant with amino acids 7-16 deleted (Hxk2^Δ7-16^), which also fail to dimerize, lose glucose-regulated Hxk2 nuclear exclusion. These experiments identify the Hxk2 N-terminal region as key in regulating Hxk2 nuclear localization, but via a completely different mechanism than that proposed in earlier studies [9,46]. Importantly, in the absence of Hxk2, glucose-repression of transcription is intact; only a few transcripts show minimal changes compared to wild-type cells (Fig 12).

In response to glucose starvation, the Glc7-Reg1 phosphatase is inactive, and the Snf1 kinase is activated by phosphorylation [77,78]. However, the loss of Snf1 or its substrate Mig1 does not dramatically alter the formation of the nuclear Hxk2 pool. In contrast, loss of Reg1 completely blocks the formation of the nuclear Hxk2 pool in glucose-starved cells. Whether Glc7-Reg1 directly dephosphorylates Hxk2 or another factor is responsible for this regulation remains to be determined.

Similarly, the Tda1 kinase is also required for starvation-induced Hxk2 nuclear accumulation. When glucose is limiting, Tda1 protein levels rise substantially, not because of altered transcription but due to increased protein stability or elevated translation. Tda1 is phosphorylated in these conditions, which activates its kinase activity towards histone substrates [62]. In glucose-starvation conditions, Tda1 phosphorylates Hxk2 at S15, as demonstrated in earlier studies, and this modification disrupts the dimer in favor of monomeric Hxk2 [35,61]. This S15 phosphorylation is likely the predominate role that Tda1 plays in glucose-starved Hxk2 regulation, given that the S15D mutation restores Hxk2 nuclear accumulation in *tda1*Δ cells.

Notably, a second layer of glucose-induced regulation must exist for Hxk2 because Hxk2^S15D^ is nuclear excluded even in glucose-grown cells. Deleting the N-terminal amino acids 7-16 or changing K13 to alanine bypasses this glucose regulation. Nuclear Hxk2 accumulation in glucose-starvation conditions is not needed for transcriptional changes, as there were little to no differences in gene expression between wild-type and *hxk2*Δ cells. Given that Hxk2 does not substantially impact global gene expression, future studies must determine its nuclear function in *S. cerevisiae*.

### Regulation of hexokinase nuclear localization in mammals

In mammals and plants, most isoforms of hexo- and glucokinases translocate to the nucleus. However, in many instances, the nuclear function is poorly understood both biochemically and mechanistically [70]. Glucokinase (GCK) regulation by glucokinase regulatory protein (GKRP) is one interesting example of nuclear regulation in mammals [79]. GCK is an important regulator of glucose balance that controls glucose metabolism in the liver and pancreas and regulates insulin secretion from β-islet cells [79–81]. In response to glucose starvation, GCK in the liver binds to GKRP which competitively inhibits glucose binding [82]. When bound to GKRP, GCK translocates into the nucleus, where it serves as a reservoir of inactive GCK that can be rapidly remobilized to the cytosol when glucose becomes available [79,82].

No yeast equivalent of GKRP has been identified, and it is unclear if Hxk2 nuclear accumulation could similarly act as a reservoir for Hxk2 function. However, it seems unlikely that nuclear Hxk2 would follow this “nuclear storage model” because even after prolonged starvation, a robust amount of Hxk2 remains in the yeast cytoplasm (S2D Fig). In addition, Hxk1 and Glk1 are both expressed in response to glucose starvation. These alternative enzymes are capable of phosphorylating glucose under starvation conditions, negating the need for a rapid return of Hxk2 to the cytosol.

Human HKII, the isoform with the highest sequence homology to Hxk1 and Hxk2, localizes to the mitochondria in glucose-replete conditions but then translocates to the nucleus in some cancer cells [27,67,69,83]. In one case, this HKII nuclear translocation is associated with apoptosis-inducing factor (AIF) and phosphorylated p53, a tumor suppressor [69,84]. When HKII moves together with these factors from the mitochondria to the nucleus, it triggers apoptosis. In yeast, Aif1 is the yeast ortholog of mammalian AIF, and loss of Hxk2 is suggested to induce apoptosis via an Aif1-dependent mechanism [85,86]. However, it is unclear if, like the mammalian model, Hxk2 is involved in the nuclear transition of Aif1 in yeast.

Others have found ∼10% of Hxk2 activity is mitochondrially associated in yeast [87], which suggests that in addition to nuclear Hxk2 there may be mitochondrially localized Hxk2. Mass spectroscopy aimed at identifying the mitochondrial proteome under glucose-replete conditions and in alternative carbon sources further supports the presence of a mitochondrial pool of Hxk1 and Hxk2 [88]. We did not observe a distinct mitochondrial pool of Hxk2 in our imaging, but it could have been masked by the bright cytosolic Hxk2. Perhaps a preexisting mitochondrial Hxk2 gives rise to nuclear Hxk2 in glucose-starvation conditions, as can be the case for mammalian HKII [85,86].

In future work, it will be critical to consider these existing models for hexokinase and glucokinase nuclear regulation and function when examining the role of nuclear Hxk2 in yeast. We are hopeful that this new paradigm for Hxk2 nuclear regulation, together with the powerful genetic approaches available in yeast, will help shed mechanistic light on the modes of hexokinase and glucokinase function in humans.

## Materials and Methods

### Yeast strains and growth conditions

Yeast strains used in this study are summarized in Supplemental Table 1 and are typically derived from the BY4742 background of *S. cerevisiae.* Where indicated, yeast cells were grown in synthetic complete (SC) medium lacking the appropriate amino acid for plasmid selection [89] with ammonium sulfate as a nitrogen source, or in yeast peptone dextrose (YPD). Plasmids were introduced into yeast strains using the lithium acetate transformation method [90]. Unless otherwise indicated, yeast cells were grown at 30°C. Liquid medium was filter sterilized. For experiments where cells were shifted into low glucose medium, yeast cells were first grown to mid-log phase in SC medium with 2% glucose (high glucose medium). Next, cells were centrifuged at 3,000 x g for 3 min, and the medium was aspirated. Cells were then resuspended in 1 mL of 0.05% glucose medium (low glucose medium), centrifuged and aspirated again to remove residual medium. Cell pellets were resuspended in SC medium with 0.05% glucose (low glucose medium) and incubated for 2 hours at 30°C unless otherwise indicated.

To assess whether Hxk2 mutants were enzymatically active in yeast cells, we performed growth assays in cells lacking chromosomal copies of the *HXK1*, *HXK2*, and *GLK1* genes. Cells lacking these hexokinases fail to grow on a glucose-containing medium because they cannot phosphorylate glucose, which is required for glycolysis [91]. These *hxk1*Δ *hxk2*Δ *glk1*Δ cells were grown in SC media lacking nutrients needed for plasmid selection and containing 2% (w/v) galactose as a carbon source. Cells were transformed with plasmids expressing the genes encoding hexokinase proteins (wild type or mutant) or an empty vector. To assay hexokinase function, cells were grown overnight in SC media containing galactose and then inoculated into SC media with glucose as the carbon source. Ninety-six well plates containing 0.2 mL of media were grown at room temperature for 36 hours. Absorbance at 600 nm was determined using a Synergy 2 plate reader (BioTek).

### Yeast protein extraction, CIP treatments, and immunoblot analyses

Whole-cell protein extracts were generated using the trichloroacetic acid (TCA) method [92]. Briefly, an equal density of mid-log phase cells (typically ∼ 5×10^7^ cells) was harvested by centrifugation, washed in water, and then resuspended in water with 0.25 M sodium hydroxide and 72 mM β-mercaptoethanol. Samples were incubated on ice, and proteins precipitated by adding 50% TCA. After further incubation on ice, proteins were collected as a pellet by centrifugation, the supernatant was removed, and the proteins were solubilized in 50 μL of TCA sample buffer (40 mM Tris-Cl [pH 8.0], 0.1 mM EDTA, 8 M Urea, 5% SDS, 1% β-mercaptoethanol, and 0.01% bromophenol blue). Samples were next heated to 37°C for 15 min, and the insoluble material was removed by centrifugation before resolving samples by SDS-PAGE. For Figure 9G, 15 μL of cell lysate was further treated for 1 h at 37°C with 40 units of Quick <u>c</u>alf intestinal alkaline <u>p</u>hosphatase (CIP, New England Biolabs, Ipswich, MA, USA) per the manufacturer’s recommendations, or mock-treated in CIP buffer without enzyme. These samples were then precipitated using 50% TCA and solubilized in SDS/Urea sample buffer as above, before analysis via SDS-PAGE. Proteins were identified by immunoblotting with a mouse anti-GFP antibody (Santa Cruz Biotechnology, Santa Cruz, CA, USA) to detect <u>g</u>reen fluorescent <u>p</u>rotein (GFP) fused to Hxk2, or mouse anti-<u>mN</u>eon <u>G</u>reen (mNG) nanobody (ChromoTek, Planegg-Martinsried, Germany) to detect mNG fused to Tda1. Primary antibodies were detected using anti-mouse or anti-rabbit secondary antibodies conjugated to IRDye-800 or IRDye-680 (Li-Cor Biosciences). Revert (Li-Cor Biosciences, Lincoln, NE, USA) total protein stain was used as a loading and membrane-transfer control. Secondary antibodies or Revert staining were detected on an Odyssey CLx infrared imaging system (Li-Cor Biosciences).

### Co-immunoprecipitation assays

Hxk2 proteins were tagged at the C-terminus with either three copies of the V5 epitope [93] or the GFP protein. Protein expression was monitored by immunoblotting with a 1:1000 dilution of Anti V5 Antibody (Fisher Scientific Catalog # R960-25) or a 1:1000 dilution of Anti-GFP polyclonal Antibody (Product # PA1-980A). Hxk2-3V5 and associated proteins were immunoprecipitated from glass bead extracts (250 μg total protein) using 20 μL of agarose conjugated anti-V5 antibody (Sigma; A7345). The extract and beads were incubated at 4°C overnight, then washed three times in 1 mL of hexokinase buffer with protease inhibitors and eluted in 15 μL SDS sample buffer.

### Recombinant protein purification and size exclusion chromatography

Hxk2 was cloned into the bacterial expression plasmid pET14b (Novagen) such that the C-terminus contained a 6-histidine tag. Hxk2 expression was induced with 1 mM IPTG at 25 °C for 2 hours. Recombinant proteins were purified on Ni-NTA columns (1 ml Ni-NTA Agarose; Qiagen Cat # 30210), washed with 20 ml hexokinase buffer (20 mM HEPES, 5 mM Mg Acetate, 100 mM NaCl, 0.5 mM EDTA, 0.5 mM DTT) with 100 mM imidazole before elution in hexokinase buffer with 500 mM imidazole. Proteins were dialyzed into hexokinase buffer with 5% glycerol and stored at −80°C.

Purified proteins were analyzed by size exclusion chromatography using a TOSOH G2000 SW_XL_ column on a Shimadzu HPLC system. Samples (40 μg protein in 50 μl of buffer) were resolved in hexokinase buffer with or without 2 mM glucose at a flow rate of 1 mL/min.

### Fluorescence microscopy

Unless otherwise indicated, imaging was performed using a live-cell microscopy protocol, maintaining yeast in the same medium they were grown in throughout the imaging process. This is a critical facet of this work because earlier studies of Hxk2 localization often fixed cells (perhaps to allow better DAPI nuclear staining) or incubated them in medium that lacked glucose and therefore likely altered Hxk2 localization. Fluorescent protein localization was performed by growing cells in SC medium with 2% glucose overnight, re-inoculating cultures at an optical density (OD)_600_ of 0.3 into fresh SC medium with 2% glucose, and growing cells until they reached mid-logarithmic phase (usually for an additional 4-5 h) at 30°C with aeration on a roller drum. For treatments in low glucose medium (SC with 0.05% glucose), cells were washed and incubated as described above in the “Yeast Strains and Growth Conditions” section. For imaging, cells were first plated on a 35 mm glass bottom microwell dish that had been coated with 15 μL (0.2 mg/mL) of concanavalin A (MatTek Corporation, Ashland, MA). They were then imaged using a Nikon Eclipse Ti2 A1R inverted confocal microscope (Nikon, Chiyoda, Tokyo, Japan) outfitted with a 100 x oil immersion objective (NA 1.49). Images were captured using GaAsP or multi-alkali photomultiplier tube detectors, and the acquisition was controlled using NIS-Elements software (Nikon). All images within an experiment were acquired using identical settings, and images were adjusted evenly and cropped using NIS-Elements.

For S1F Figure, cells were grown as described [11]. 25 μL of cells were loaded onto ConA-coated glass slides. Cells were allowed to settle for a couple of minutes. Then the remaining suspension was aspirated off the slide. DAPI (1 μL of 2.5 μg/mL dissolved in 80% glycerol) was added to the cells. A glass coverslip was placed over the cells [11] and then imaged as described above, this time using a Nikon Eclipse Ti2-E inverted microscope (Nikon, Chiyoda, Tokyo, Japan) equipped with an Apo100X objective (NA 1.45) and captured with an Orca Flash 4.0 cMOS (Hamamatsu, Bridgewater, NJ) camera and NIS-elements software (Nikon). These conditions mirror those used in earlier publications that assessed Hxk2-GFP localization [9–11,36,46].

Fluorescence recovery after photobleaching (FRAP) experiments (see Fig 2E, S2 Fig, and S1-2 Movies) were performed by adding 25 μL of low-glucose incubated cells to ConA-coated Matek dishes. The experiment was conducted using the confocal microscope described above. First, images were captured before the nuclei were bleached. Next, a region of interest (ROI) for bleaching (see Su2E Fig) was defined in the cell nuclei, using the mScarlet nuclear marker as reference. A 1 sec pulse of the 488 nm laser (10% power) was used to bleach the nuclei. An image of the bleached cells was captured immediately after and then every minute for 20 mins to monitor nuclear fluorescence recovery.

### Image quantification and statistical analyses

Quantification of nuclear fluorescence and whole-cell fluorescence intensity was done using Nikon General Analysis 3 software (Nikon, Chiyoda, Tokyo, Japan) with the segmentation from NIS-Elements.*ai* (Artificial Intelligence) software (Nikon, Chiyoda, Tokyo, Japan) unless otherwise described. For quantification of whole-cell fluorescence, the NIS.*ai* software was trained on a ground truth set of samples where cells had been manually segmented using the DIC channel images. Next, the NIS.*ai* software was iteratively trained until it achieved a training loss threshold of <0.02, indicating a high degree of agreement between the initial ground truth and the output generated by the NIS.*ai* software. To measure the mean nuclear fluorescence, we trained the NIS.*ai* software to identify the nucleus using the chromosomally integrated Tpa1-mScarlet nuclear marker (see S2 Table). The NIS.*ai* software was trained using a manually defined ground truth set of nuclear segmentations. Using the General Analysis 3 software, fields of images captured through confocal microscopy were processed so that individual whole-cell and nuclear objects in a field of view were segmented using the DIC and 561 nm (mScarlet) channels. A parent-child relationship was applied to individual nuclear objects (child) within the same cell (parent) to aggregate them as single objects and pair them to the appropriate whole cell. Any partial cells at the edges of the image were removed along with their child objects. Then the mean fluorescence intensity of each parent or child object was defined in the appropriate channel. All imaging quantification graphs, except for manual quantification data in S1A-B Fig and S1D-E Fig, were derived using this method.

Manual quantification to measure mean nuclear or whole-cell fluorescence (Fig 1A-B and 1D-E) was performed using ImageJ software (National Institutes of Health, Bethesda, MD). A 2-pixel wide line was hand drawn around the nuclei using images of the Tpa1-mScarlet nuclear marker to create a mask that was then overlaid on the GFP images, and the mean GFP signal was measured. The same was done to measure whole-cell fluorescence, except lines were drawn around the perimeter of each cell using the GFP or mNG signal since Hxk2 has a diffuse cytosolic distribution. Mean background fluorescence intensity was measured for each image and subtracted from the mean fluorescence measurements to calculate mean nuclear and whole-cell intensities.

Statistical analyses of fluorescence quantification was done using Prism (GraphPad Software, San Diego, CA). Unless otherwise indicated, we performed Kruskal-Wallis statistical tests with Dunn’s posthoc correction for multiple comparisons. In all cases, significant p-values from these tests are represented as: * p-value<0.1; ** p-value<0.01; *** p-value<0.001; **** p-value<0.0001; not significant (ns) = p-value>0.1. In some instances where multiple comparisons are made, the † symbol may be used in place of * to indicate the same p-values but relative to a different reference sample (see the figure legends).

FRAP data were analyzed first by measuring the mean nuclear fluorescence of a nuclear ROI both before and after bleaching. In some cases, nuclei shifted positions at different time points along the lateral plane. In these cases, we manually re-positioned the ROI and re-measured to ensure we did not erroneously measure the cytosolic pool. To account for non-specific photobleaching that occurred due to repeated rounds of imaging, an ROI reference was used in an adjacent cell when no targeted lase bleaching was performed. The mean cytosolic GFP fluorescence in the ROI of the control cell was measured at each time point as well. Then the data were normalized using the following equation [94]:

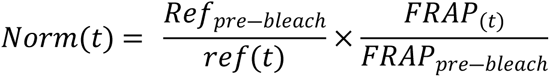

The normalized data were plotted over time, and the rates of recovery were calculated by measuring the slope of the linear portion of each recovery plot.

### RNA-Seq sample preparation and analyses

RNA samples were prepared from multiple independent yeast cultures grown on synthetic complete medium using the RNeasy Mini Kit (Qiagen). Sequencing libraries were prepared using the TruSeq Stranded mRNA library method (Illumina). RNA sequences were mapped to *S. cerevisiae* mRNA using the kallisto software package [95]. Each RNA sample yielded 40–50 million reads. mRNA abundance was expressed in transcripts per million mapped reads (tpm). To compare mRNA expression under different conditions, we used a Student’s t-test to calculate p values with a false discovery rate threshold of 0.01%. All RNA-seq data have been deposited in the SRA database under accession number PRJNA885127.

### Homology models of the *Sc*Hxk2 dimer

We generated a homology model of the *Sc*Hxk2 dimer using SWISS-MODEL [96–100]. A *Kl*Hxk1 crystallographic dimer (PDB ID 3O1W [43]) served as the template. We removed all alternate locations from the 3O1W PDB file so that each amino acid had only one sidechain conformation. We then copied the two chains, A and B, into two separate files, each containing the respective *Kl*Hxk1 monomer. We separately uploaded these two monomers to the SWISS-MODEL server, together with the full-length sequence of *Sc*Hxk2.

Since the 3O1W template structure covers *Kl*Hxk1 almost entirely—including the critical N-terminal tails—the homology models of each monomer included all *Sc*Hxk2 amino acids except the initial methionine and the terminal alanine. The initial methionine (M1) is cleaved *in vivo* [41,101], so we used UCSF Chimera [102] to add only the C-terminal alanine. To merge the two monomers into a single dimer model, we used multiseq [103], as implemented in VMD [104], to align each monomer to its respective chain in the original 3O1W dimeric structure. Finally, we processed the dimer model with PDB2PQR [105,106], which added hydrogen atoms per the PROPKA algorithm (pH 7.00) [107] and optimized the hydrogen-bond network.

To generate a final model of the *apo* (ligand-free) *Sc*Hxk2 dimer, we subjected the dimer homology model to one round of minimization in Schrödinger Maestro. To generate a final model of the *holo* (glucose-bound) *Sc*Hxk2 dimer, we aligned a glucose-bound *Kl*Hxk2 dimer (PDB ID 3O5B [43]) to our *Sc*Hxk2 dimer model and copied the aligned glucose molecules. We then added hydrogen atoms to the glucose molecules using Schrödinger Maestro. To resolve minor steric clashes and optimize interactions between the receptor and glucose ligands, we minimized the *Sc*Hxk2/glucose complex using a stepwise protocol. First, we used Schrödinger Maestro to subject all binding-site atoms (excluding the ligand) to one round of minimization. Second, we subjected all protein atoms to two rounds of minimization. Third, we subjected all protein and glucose hydrogen atoms to one round of minimization. Finally, we minimized all the atoms of the complex.

To generate a model of the *holo* (glucose-bound) *Sc*Hxk2 monomer, we simply deleted one of the monomers of our *holo Sc*Hxk2 dimer model. We note that the ATP depicted in Fig 3G and 4A was not part of the model itself (i.e., it was not included in the minimization procedure). We positioned ATP in the active site for visualization purposes by aligning a crystal structure of a homologous protein (6PDT [38]) to each monomer of our *Sc*Hxk2 dimer model.

### Molecular dynamics simulations

We performed molecular dynamics (MD) simulations of the *apo* dimer, *holo* dimer, and *holo* monomer systems. We used *tleap* (Ambertools18 [108]) in each case to add a water box extending 10 Å beyond the protein along all dimensions. We also added Na+ counter ions sufficient to achieve electrical neutrality and then additional Na+ and Cl-counter ions to approximate a 150 mM solution. The protein, counter ions, water molecules, and glucose molecules were parameterized per the Amber ff14SB [109], TIP3P [110], and GLYCAM_06j-1 [108] force fields, respectively.

We applied four rounds of minimization using the Amber MD engine [111,112]. First, we minimized all hydrogen atoms for 5,000 steps. Second, we minimized all hydrogen atoms and water molecules for 5,000 steps. Third, we minimized all hydrogen atoms, water molecules, and protein side chains for 5,000 steps. Finally, we minimized all atoms for 10,000 steps.

After minimization, we equilibrated each system using three rounds of simulation. First, we subjected each system to a short simulation in the canonical ensemble (NVT, 0.02 ns total), with a 1.0 kcal/mol/Å^2^ restraining force applied to the backbone atoms. Using the same backbone restraints, we continued the simulation in the isothermal– isobaric ensemble (NPT, 1 atm, 1.0 ns total). Finally, we finished the equilibration (NPT, 1 atm, 1.0 ns total) without restraints. In all cases, we used a 2-fs timestep and a 310 K temperature setting.

After minimization and equilibration, we ran three isothermal-isobaric (NPT, 310 K, 1 atm) productive simulations of the *Sc*Hxk2 *apo* dimer (550 ns, 262 ns, 266 ns), the *Sc*Hxk2 *holo* dimer (1000 ns, 261 ns, 262 ns), and the *Sc*Hxk2 *holo* monomer (530 ns, 253 ns, 260 ns). We used a different random seed for each simulation.

### RMS and pairwise distance analyses

To calculate how far a bound glucose molecule deviated from its initial position, we used VMD [104] to align the associated *Sc*Hxk2 monomer by its alpha carbons. We then calculated the heavy-atom root mean square distances (RMSDs) between the starting glucose pose and the pose of each frame.

To monitor the hydrogen bond between K13 and Q142*, we used VMD to calculate the distance between the K13 terminal nitrogen atom and the Q142* sidechain carbonyl oxygen atom. We assumed a hydrogen bond had formed when this distance was less than 4.0 Å. To monitor the salt bridge between K13 and D106*, we calculated the distance between the K13 terminal nitrogen atom and the D106* terminal-most carbon atom. We assumed a salt bridge had formed when this distance was less than 4.0 Å.

### Molecular visualization

Images were generated using VMD [104] and BlendMol [113].

## Acknowledgments

We acknowledge the support of the Dietrich School of Arts and Sciences Microscopy and Imaging Suite (RRID: SCR_022084). We gratefully acknowledge the support of Dr. Patrick Thibodeau for help with our size-exclusion chromatography assays. We kindly acknowledge the feedback provided by Dr. Jeff Brodsky and his research team as well as the Pittsburgh Area Yeast Meeting members prior to submission. We further acknowledge the support and feedback from members of the O’Donnell lab team.

